# A transcriptional toolbox for exploring peripheral neuro-immune interactions

**DOI:** 10.1101/813980

**Authors:** Zhi Liang, Zoe Hore, Peter Harley, Federico Uchenna Stanley, Aleksandra Michrowska, Monica Dahiya, Federica La Russa, Sara E. Jager, Sara Villa-Hernandez, Franziska Denk

## Abstract

Correct communication between immune cells and peripheral neurons is crucial for the protection of our bodies. Its breakdown is observed in many common, often painful conditions, including arthritis, neuropathies and inflammatory bowel or bladder disease. Here, we have characterised the immune response in a mouse model of neuropathic pain using flow cytometry and cell-type specific RNA sequencing (RNA-seq). We found few striking sex differences, but a very persistent inflammatory response, with increased numbers of monocytes and macrophages up to 3½ months after the initial injury. This raises the question of whether the commonly used categorisation of pain into “inflammatory” and “neuropathic” is one that is mechanistically appropriate. Finally, we collated our data with other published RNA-seq datasets on neurons, macrophages and Schwann cells in naïve and nerve injury states. The result is a practical web-based tool for the transcriptional data-mining of peripheral neuroimmune interactions.

http://rna-seq-browser.herokuapp.com/

## Introduction

The immune and peripheral nervous systems are closely entwined. A complex network of sensory and sympathetic fibres innervates our organs and engages in bi-directional communication with a host of tissue resident immune cells, including macrophages, dendritic cells and T lymphocytes. A disruption in this neuro-immune connection contributes to many common debilitating conditions, including irritable bowel disorders, diabetes, psoriasis, asthma, and chronic pain, collectively accounting for nearly 30% of all years lost to disability world-wide^1^. This article is particularly focused on pain, which in its acute form is a natural consequence and cardinal sign of inflammation^2^. In contrast, chronic pain can become uncoupled from underlying disease, causing independent nervous system dysfunction which requires bespoke treatment approaches. In recognition of this, chronic pain syndromes have just been added to the World Health Organisation’s International Classification of Diseases (ICD-11)^3^.

A primary driver for the uncoupling between pain and acute inflammation is believed to be release of pro-algesic mediators from non-neuronal cell types. This process, known as peripheral sensitisation, causes sensory neurons to become ectopically active and hypersensitive to normally innocuous stimuli^4–6^. Many different cell types have been implicated in peripheral sensitisation, including immune cells, satellite glial cells and Schwann cells^7–9^. Already two decades ago, for example, both neutrophils and macrophages were reported to be causally involved in hyperalgesia - an exaggerated pain sensation in response to a noxious stimulus^10, 11^. Nevertheless, the details of this connection remain vague, with agreement in the pre-clinical literature largely limited to the one main finding that interfering with various sub-types of myeloid cell will alter evoked pain behaviour in various animal models of persistent pain^12–15^. Mechanistically, this has been ascribed to different mediators putatively released from macrophages and neutrophils, chiefly among them TNF, IL-1β, IL-6 and NGF^7, 8, 16^.

The current lack of precision is hardly surprising when we consider the incredible cellular and functional complexity of our bodies’ inflammatory process^17^. Each tissue mounts its own distinct local inflammatory responses that, at their peak, are usually accompanied by systemic endocrine effects resulting in diverse symptoms including fever and fatigue. The correct interplay of these responses is essential for efficient immunity and tissue healing, as is a very tightly controlled resolution process – which again relies on several active, parallel inter- and intra-cellular mechanisms^18, 19^.

Here, we aim to provide a more detailed view of immune cell responses in a persistent pain state. We used partial sciatic nerve ligation as a model of traumatic neuropathic pain and performed flow cytometry analysis and fluorescence-activated cell sorting on male and female mice over a 3½ month time course. RNA-seq was performed at 1 week and 10 weeks following nerve injury on immune cell sub-populations (Figure 1A) extracted from sciatic nerve and the ganglia in which its cell bodies are located: lumbar L3-L5 dorsal root ganglia (DRG). Our results were integrated with other publicly available data of relevant cell types to generate a plug-and-play, user-friendly website for interrogating peripheral neuro-immune interactions at transcriptional level.

**Figure 1:**
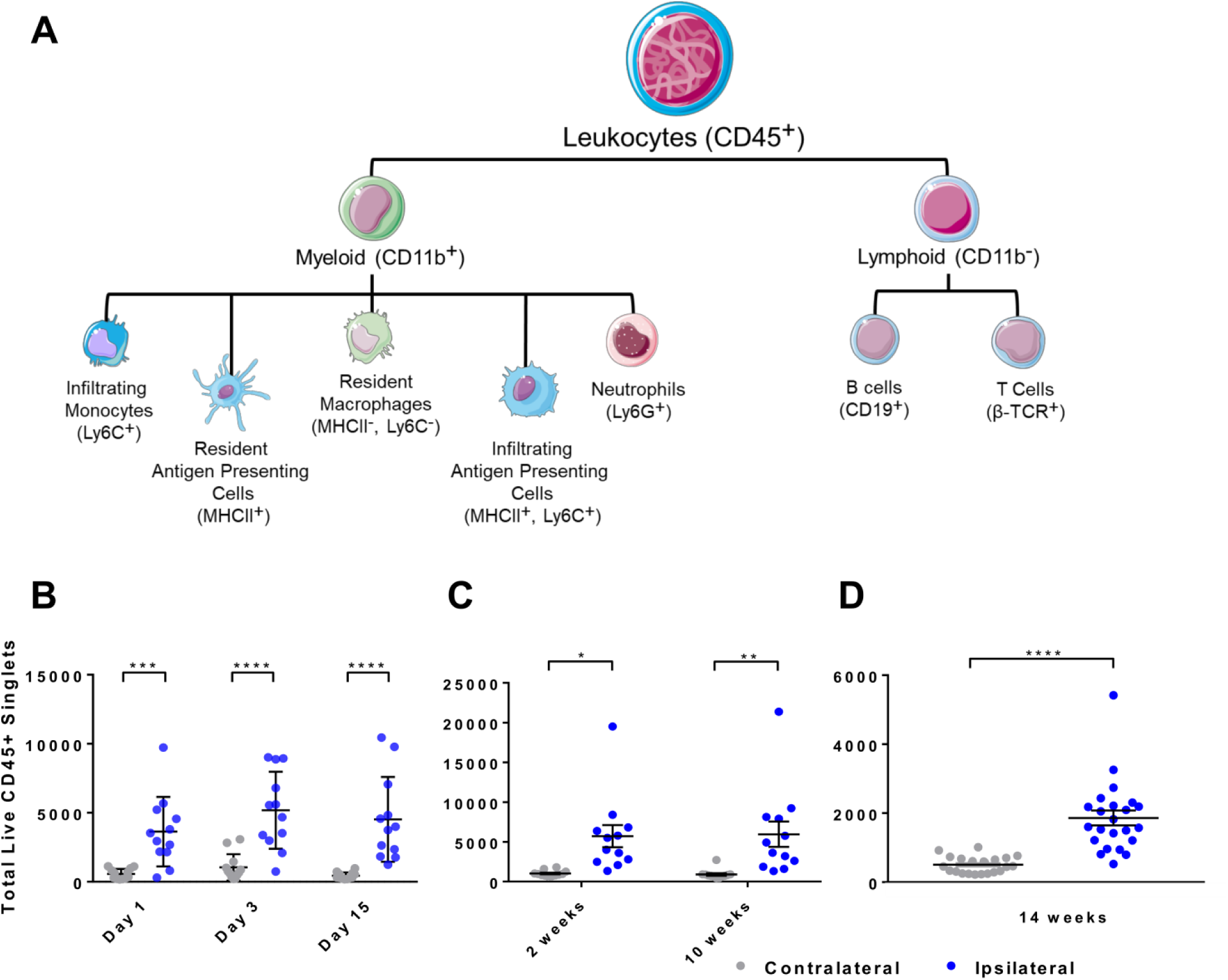
Partial sciatic nerve ligation induces a long-lasting increase in total CD45+ immune cell counts in sciatic nerve. **A**, Schematic of the markers used to differentiate lymphoid and myeloid immune cell populations in flow cytometry and FACS experiments. **B-D**, Plotted are the total number of live CD45+ singlets isolated from sciatic nerve after partial sciatic nerve ligation at different time points. Results were obtained as part of three separate experimental batches 8-D. In **B & C**, n = 12 mice per time point (6 males & 6 females) were processed, whilst in **D**, n = 23 mice (11 males & 12 females) were processed, shown here as separate data points with mean ±SEM.

## Results

In order to develop a detailed characterisation of immune responses in the mouse sciatic nerve following partial sciatic nerve ligation (PSNL), we performed three batches of flow cytometry experiments at different time points. In each instance, we used a minimum of 6 male and 6 female mice. In every experiment, the total number of live CD45+ immune cells was significantly elevated in ipsilateral nerves compared to their contralateral counterparts (Figure 1B-D). For instance, 14 weeks (3½ months) after PSNL, there were on average 3.7x more immune cells in ipsilateral nerve, a highly statistically significant result (Welch’s independent samples t-test, t(24)=6.13, p < 0.0001). A direct, batch-controlled comparison of immune cells collected 2 weeks and 10 weeks (2½ months) after PSNL showed that there was no major difference in immune cell numbers at these two points, with a two-way ANOVA revealing no significant main effect (F(1,44)=0.01, p = 0.96) or interaction with time (F(1,44)=0.03, p=0.87). The effect of injury on cell number was once more highly significant: F(1,44)=20.93, p < 0.0001.

This is quite striking given that, macroscopically, the site of injury is drastically different weeks versus months after ligation (Suppl. Figure 1): 2-3 months after surgery, any visible signs of the suture have completely disappeared, with only slight swelling and discoloration remaining on the ipsilateral side. Nevertheless, mice still display overt pain-like behaviours at 3½ months, with paws being visibly clenched and/or placed gingerly on any surface (Suppl. Figure 2). Moreover, using immunofluorescence, high numbers of immune cells were still clearly visible 14 weeks after PSNL (Suppl. Figure 3).

Detailed flow cytometry analysis revealed that the persistent increase in CD45+ cells is due to an increase in all immune cell types present in nerve, with the exception of Ly6G+ neutrophils (Figures 2, and Suppl. Figure 4). We observed MHCII+ single positive myeloid cells and MHCII-/LyC6-double negative myeloid cells in contralateral nerves. After nerve injury, two additional populations appeared: Ly6C+ single positive and MHCII+/Ly6C+ double positive cells (see Suppl. Figures 5 and 6 for gating strategies, and Suppl. Table 1 for panels). All 4 populations were significantly upregulated 2 weeks, 10 weeks and 14 weeks after injury. Comparing 2 weeks vs. 10 weeks, there was no significant main effect of time (F(1,22)=0.48, p = 0.495) nor an interaction between time and injury (F(1,22)=0.33, p = 0.572).

**Figure 2:**
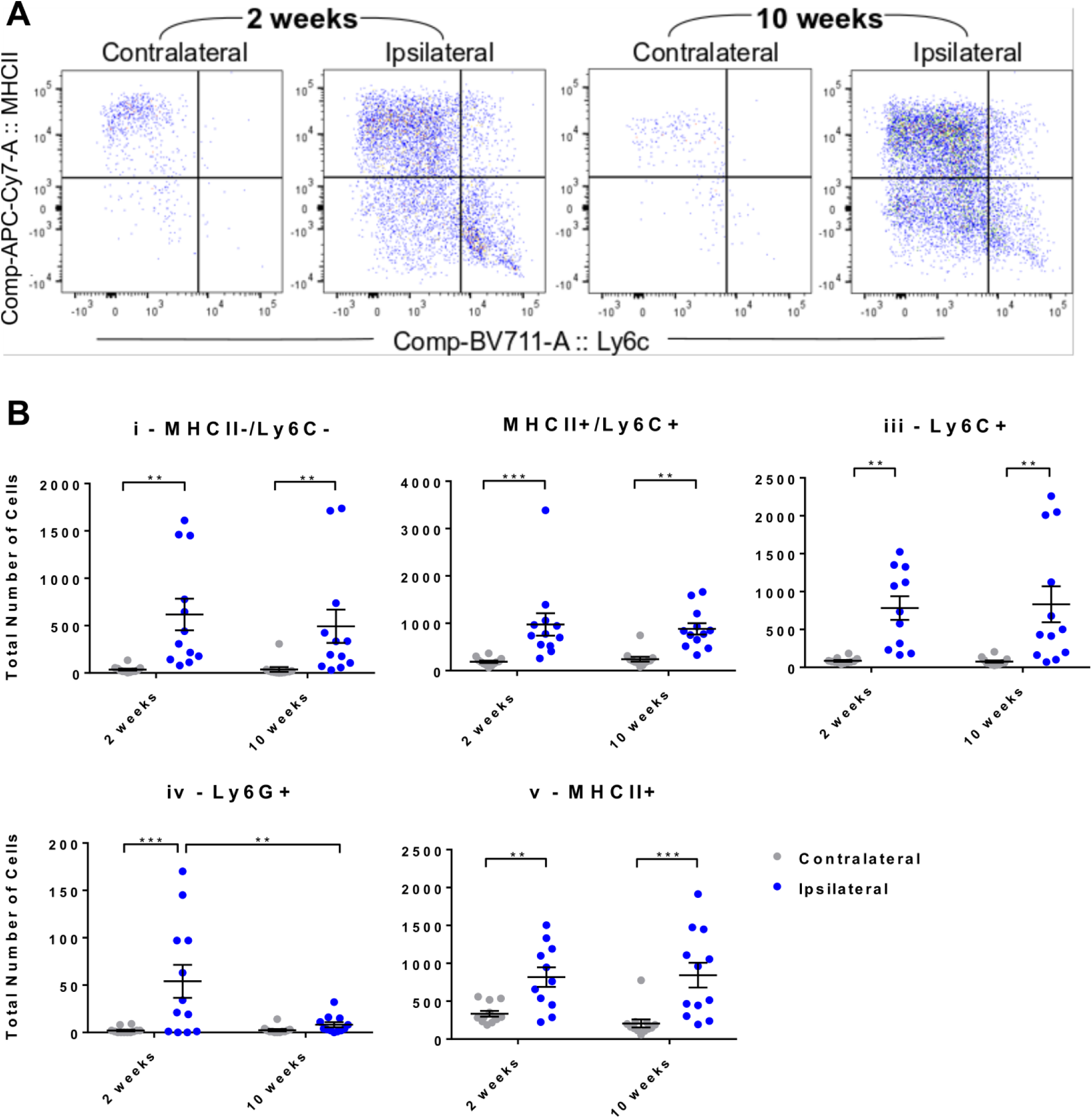
All major myeloid cell populations apart from neutrophils are still upregulated in sciatic nerve two and a half months after partial nerve ligation. **A)** Flow cytometry was used to phenotype the major myeloid cell populations in sciatic nerve. Shown here are representative dot plots of four of the five major myeloid subpopulations, gated on live CD45+, CD11b+, Ly6G-singlets: MHCII+ (putative resident, antigen presenting macrophages), MHCII+/Ly6C+ (likely infiltrating monocytes differentiating into resident populations), Ly6C+ (infiltrating monocytes), MHCII-/Ly6C-(likely resident macrophages). Note the presence of the MHCII+/Ly6C+ double positive population and infiltrating Ly6C+ monocytes (hi and Io) primarily after nerve injury. **B)** Quantification of the total number of live CD45+/CD11b+ singlets obtained from ipsilateral and contralateral nerves which were either negative for **i)** MHCII-/Ly6C- or positive for **ii)** MHCII+/Ly6C+, **iii)** Ly6C+, **iv)** Ly6G+ (neutrophils) or **v)** MHCII+. At each time point, n = 12 mice were processed (6 males & 6 females), shown here as separate data points with mean ±SEM.

Similar to CD11b+/Ly6G-populations, αβ+ T cells and CD11b-/Ly6G-populations were also increased on the ipsilateral side (Figure 3 and Suppl. Figure 4), starting from around 3 days after injury. Absolute event counts remained elevated up to 14 weeks after injury. As we observed previously in DRG^20^, nerve seemed to contain a large population of unidentified CD45+ cells that are non-myeloid (CD11b-) and not positive for αβTCR. They also do not appear to be positive for CD19 (B cells), and preliminary trials indicated that they are negative for γδ+ TCR or glutamine synthetase, a satellite glial cell marker (data not shown). In contrast to CD11b+/Ly6G-populations, neutrophil numbers (CD11b+/Ly6G+) peaked at day 1 and were still somewhat elevated 2 weeks after PSNL, but then dropped back to contralateral levels (Figure 2 **&** Suppl. Figure 7).

**Figure 3:**
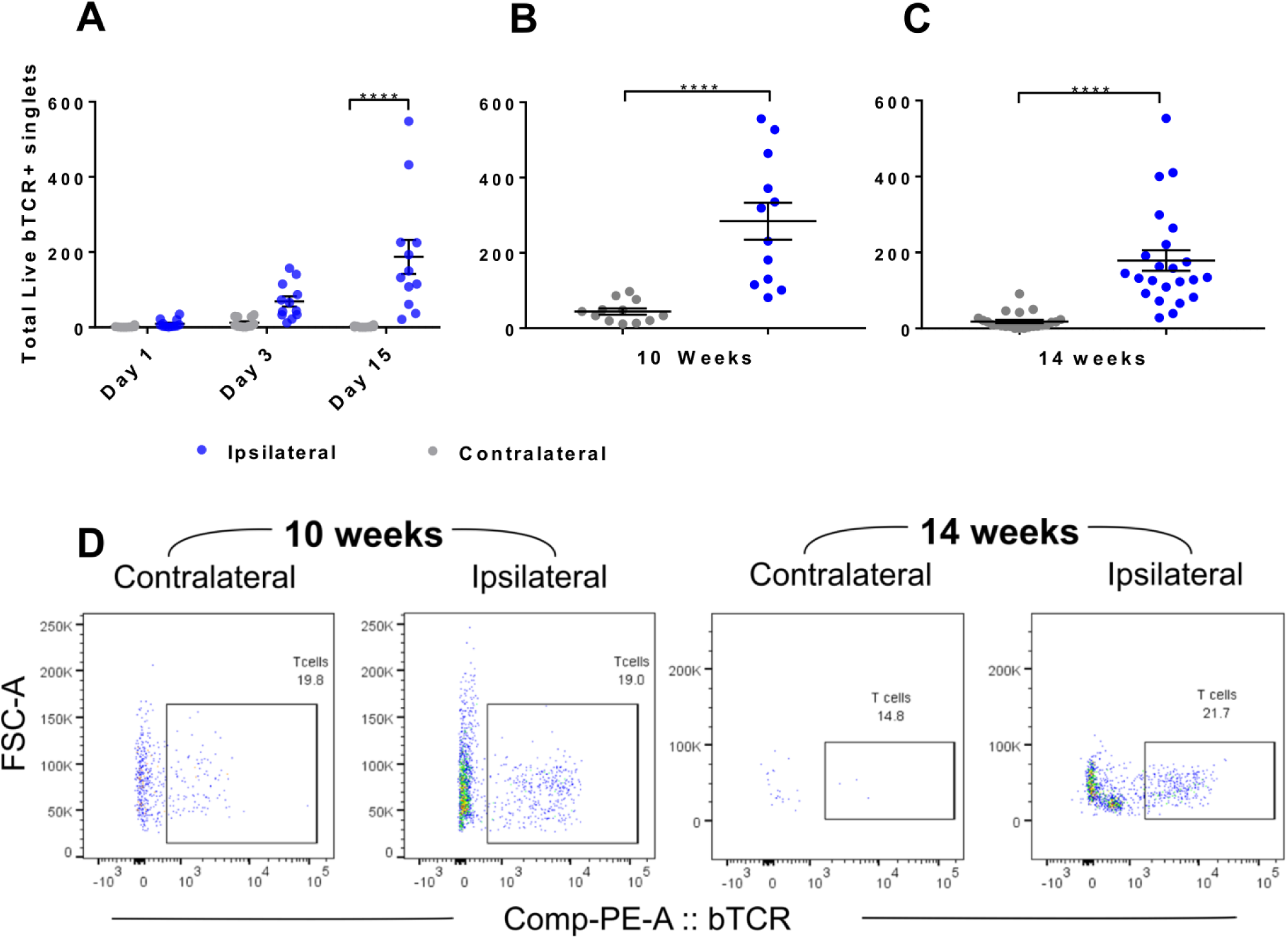
Partial sciatic nerve ligation induces a long-lasting increase in aji T cell counts in sciatic nerve. Plotted are the total number of live CD45+/CD11b-/αβTCR+ singlets isolated from sciatic nerve after partial sciatic nerve ligation at different time points. Results were obtained as part of three separate experimental batches (**A-C**). In **A & 8**, n = 12 mice per time point (6 males & 6 females) were processed, whilst in **C**, n = 23 mice (11 males & 12 females) were processed, shown here as separate data points with mean± SEM. **D**) Representative dot plot from ipsilateral and contralateral nerves at 10 weeks and 14 weeks after injury, also showing the presence of a large unknown population (CD45+/CD11b-/αβTCR-).

The picture emerging from DRG was different (Figures 4 & 5), with the increase in the total number of live CD45+ cells being much less pronounced. A two-way ANOVA revealed significant main effects of injury (F(1,43)=13.03, p < 0.001) and time (F(1,43)=5.44, p = 0.024), with a significant increase in immune cell numbers in ipsilateral DRG only at 2 weeks (adj. p = 0.003). Of note, we detected a permanent CD45+/CD11b+/Ly6G+ positive population in DRG, even on the contralateral side, indicating that Ly6G+ might not be an ideal marker for neutrophils in this tissue (Figure 5 & Suppl. Figure 8).

**Figure 4:**
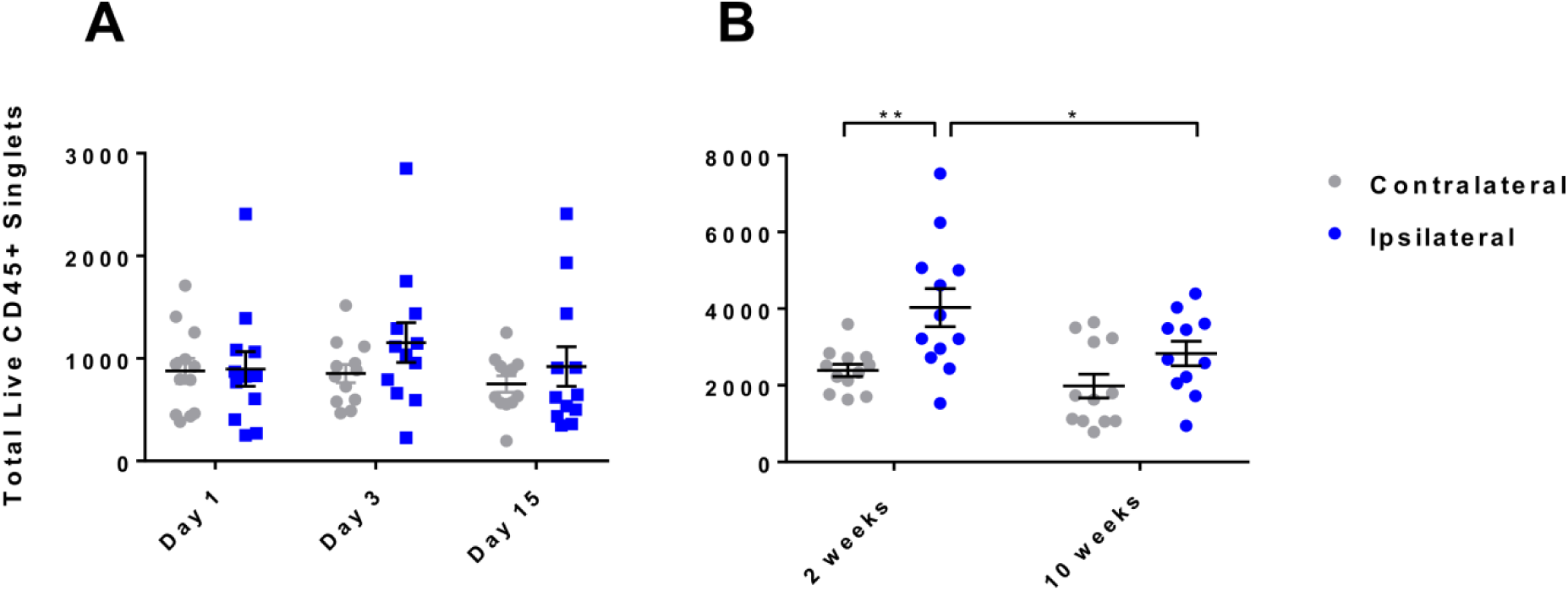
Total CD45+ immune cell counts in DRG are less variable over time and any increases do not persist long-term. Plotted are the total number of live CD45+ singlets isolated from L3-5 DRG after partial sciatic nerve ligation at different time points. Results were obtained as part of two separate experimental batches (**A & B**). In both instances, n = 12 mice per time point were processed (6 males & 6 females), shown here as separate data points with mean ±SEM.

**Figure 5:**
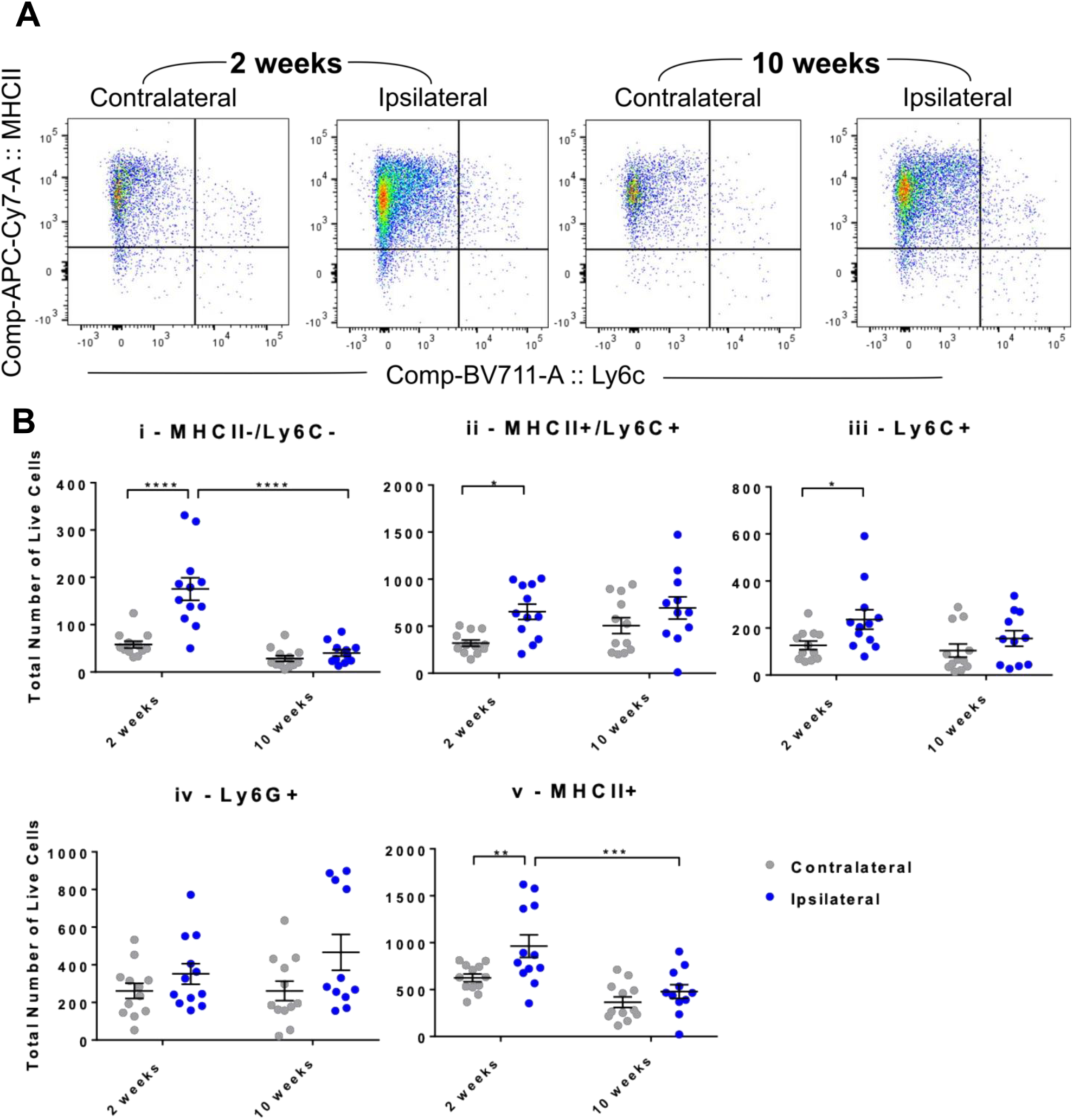
None of the major myeloid cell populations in the DRG remain significantly upregulated two and a half months following partial sciatic nerve ligation. **A)** Flow cytometry was used to phenotype immune cell populations in DRG. Shown here are representative dot plots of myeloid subpopulations, gated on live CD45+, CD11b+, Ly6G-singlets: MHCII+ (putative resident, antigen presenting macrophages), MHCII+/Ly6C+ (likely infiltrating monocytes differentiating into resident populations), Ly6C+ (infiltrating monocytes), MHCII-/Ly6C-(likely resident macrophages). **B)** Quantification of the total number of live CD45+/CD11b+ singlets obtained from ipsilateral and contralateral DRG which were either negative for **i)** MHCII-/Ly6C- or positive for **ii)** MHCII+/Ly6C+, **iii)** Ly6C+, **iv)** Ly6G+ or **v)** MHCII+. At each time point, n = 12 mice were processed (6 males & 6 females), shown here as separate data points with mean ±SEM. Note that Ly6G in the DRG does not vary with injury or time (see text).

**Figure 6:**
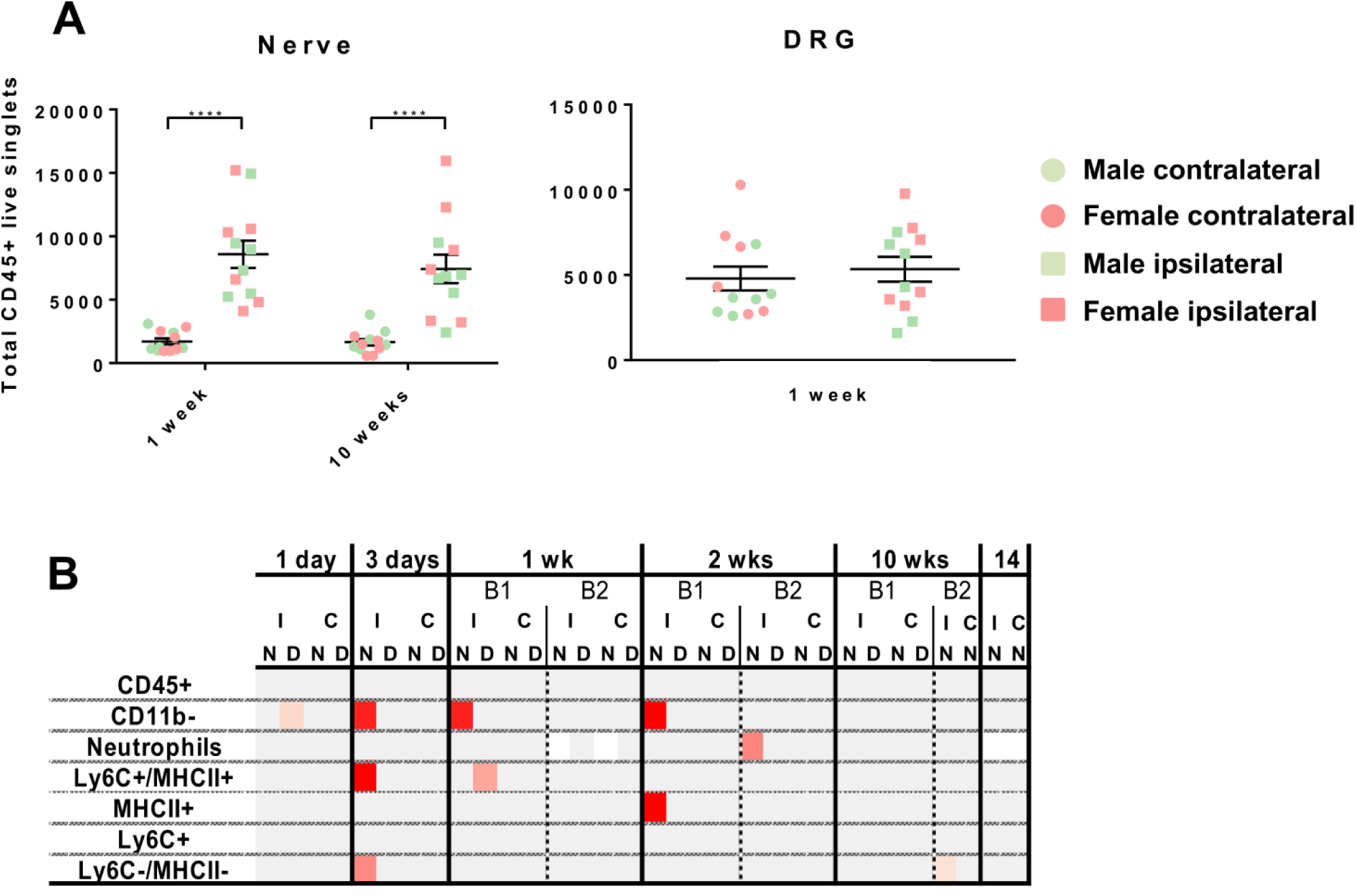
There are no striking sex differences in immune cell numbers in either nerve or DRG after partial sciatic nerve ligation at any of the time points. **A)** Plotted are select examples of the total number of live CD45+ singlets obtained from ipsilateral and contralateral nerves and DRG from male and female mice. At each time point (1 and 10 weeks after PSNL for nerve, 1 week after PSNL for DRG), n = 12 mice were processed (6 males & 6 females), shown here as separate data points with mean ±SEM. **B)** Over the course of this work, immune cells were characterized at 6 different time points, with two batches (B) each for 1 week, 2 weeks and 10 weeks after nerve injury. At each time point we had n = 6 per male/female group (with the exception of week 14, where we had n=11/12 per group). Plotted here is a summary of the sex differences we observed for all major populations from nerve (N) and DRG (D) in ipsi-(I) and contralateral (C) tissue. White squares: no data collected; grey squares: no significant sex differences; graded red squares: significant sex differences with p values ranging from p < 0.05 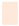 to p < 0.03 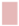 and p < 0.005 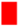. No consistent differences emerged, suggesting that any single instance might be a false-positive event.

We previously detected sex differences in lymphocyte numbers when examining immune cells from DRG from a smaller number of mice (n = 4, male vs. female) at two time points (day 8 vs. day 28)^20^. Our current, more thorough exploration, failed to replicate this. We were unable to detect any consistent differences in immune cell numbers in either nerve or DRG after partial sciatic nerve ligation at any of the time points examined (Figure 6A). Statistically significant differences did emerge in particular experimental batches, but failed to consistently replicate (Figure 6B).

Our flow cytometry results were complemented by transcriptional experiments. To gain a broad overview of different molecular phenotypes, we performed shallow sequencing on various myeloid cell subpopulations in nerve and DRG 1 and 10 weeks after PSNL, and also examined T cells from nerve at 10 weeks after PSNL. Samples included in the subsequent analyses were screened for sequencing quality: on average 95% of reads fell within genes, there was negligible ribosomal (1% on average) and no DNA contamination (**Suppl. Table 2** and **Suppl. Tables 3 & 4** for processed TPM values). On average, 7.8M of high quality, non-duplicate reads were captured. Samples were also relatively free of contaminating cell types, with the exception of DRG macrophages which suffered from a small degree of contamination with satellite glial and neuronal transcripts (Suppl. Figure 9). This should be considered when interpreting differentially expressed transcripts between nerve and DRG myeloid cells.

As expected, our sequencing samples at week 1 segregated by myeloid cell phenotype, although principle component analysis also indicated quite significant transcriptomic changes as a result of injury (Figure 7), particularly in MHCII+ myeloid cells from sciatic nerve. Indeed, 279 transcripts were identified as significantly dysregulated in ipsilateral MHCII+ cells compared to their contralateral counterparts (*sleuth* algorithm^21^, adj. p < 0.05, **Suppl. Table 5**).

**Figure 7:**
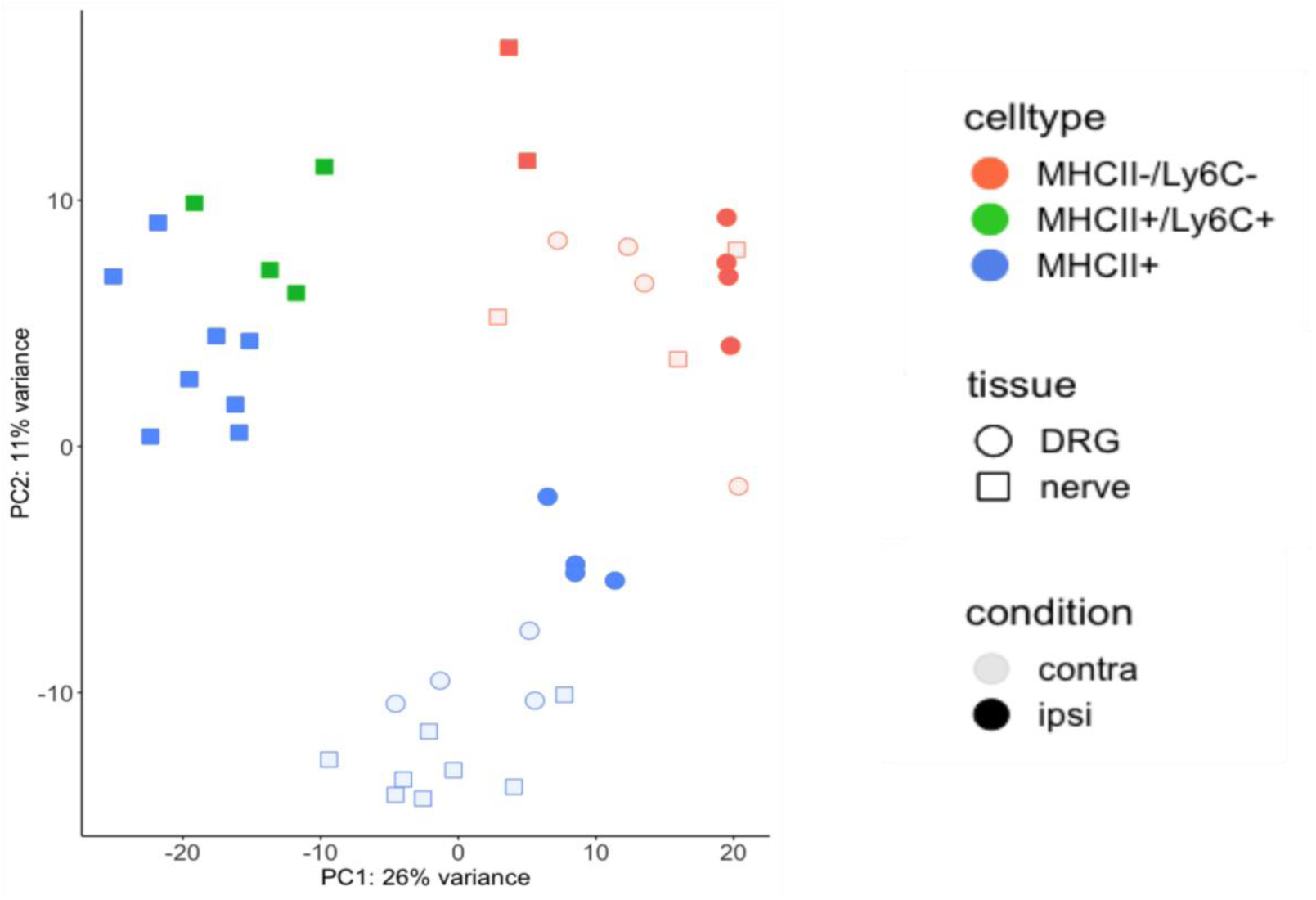
Principle component analysis (PCA) of FAC-sorted immune cells show they cluster by cell type, as expected, but also by injury state - suggesting significant transcriptomic changes. PCA of all samples extracted and sequenced 1 week after PSNL. The three main visible clusters are: 1) ipsilateral MHCII+ populations from nerve together MHCII+/Ly6C+ populations which are only present in an injury state; 2) MHCII-/Ly6C- resident populations and 3) MHCII+ contralateral nerve and ipsi- and contralateral DRG populations.

Subsequent network analysis suggested that, unsurprisingly, MHCII+ cells in injured nerve upregulate genes related to antigen presentation and lymphocyte interactions, but downregulate genes related to more canonical pro-inflammatory and resident macrophage functions (Figure 8). Thus, we for instance found significantly increased expression of several MHCII genes (H2-K1, H2Q7, H2-Oa, H2-Ob) and of Itgax, the gene encoding for the dendritic cell marker CD11c. We validated the latter result at protein level (Suppl. Figure 10), recording significantly more CD11c expressing cells amongst the MHCII+/Ly6C-population in sciatic nerve three and seven days after PSNL.

**Figure 8:**
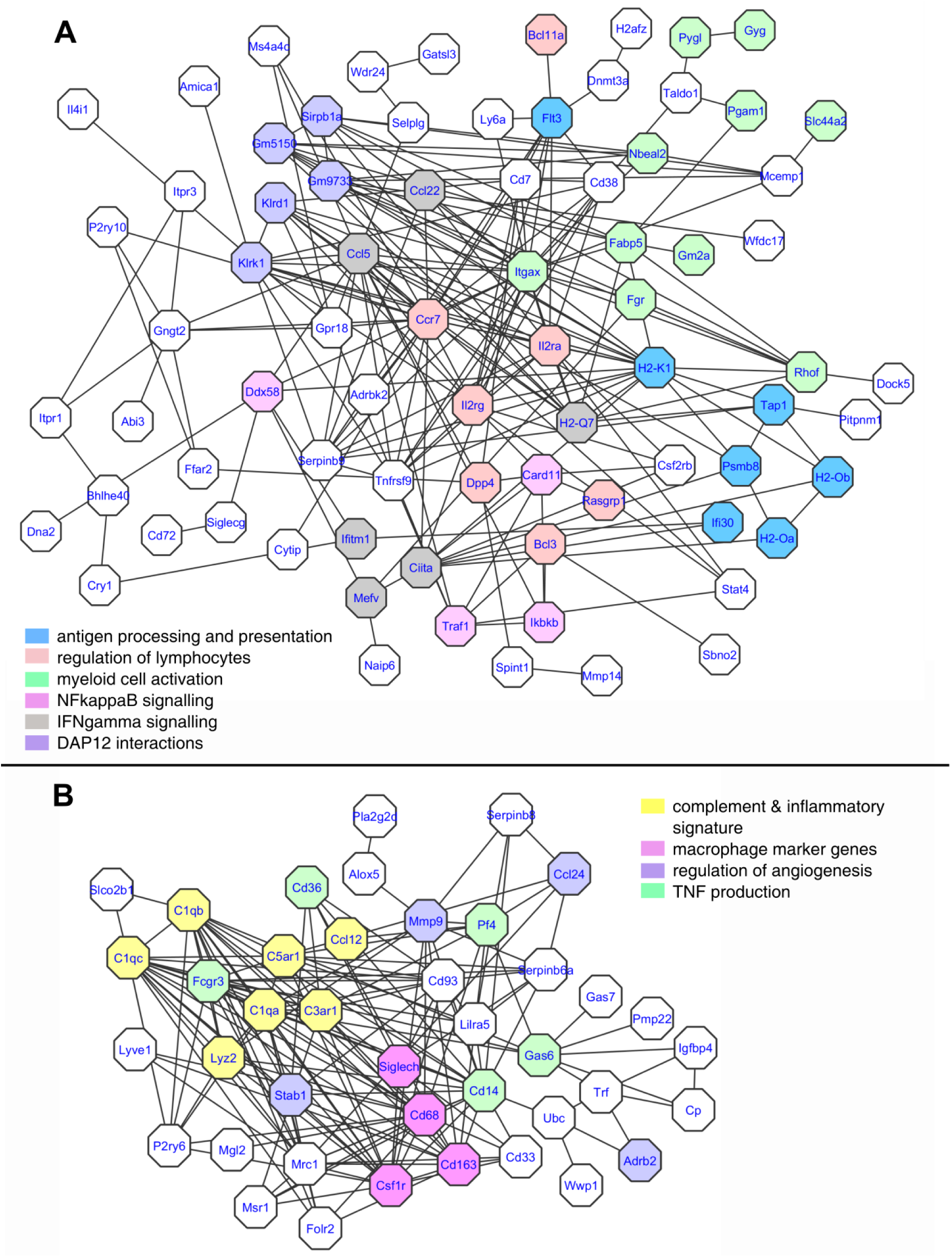
At one week post PSNL, MHCII+/Ly6C-myeloid cells from sciatic nerve upregulate functions relating to interactions with other immune cells, in favour of more generic pro­inflammatory and homeostatic activities. **A)** STRING network analysis reveals that 77 of 186 genes upregulated in ipsilateral MHCII+ macrophages at adj. p < 0.05 are likely to be functionally connected with overrepresented processes including antigen presentation, regulation of lymphocytes and myeloid cell activation. **B)** Conversely, 41 of 93 significantly downregulated genes formed a network that includes transcripts relating to pro-inflammatory function (TNF, complements), regulation of angiogenesis and canonical macrophage markers, like CD163 typically found in resident macrophages. See **Suppl. Table 5** for differential expression tables.

Also in keeping with our cell count data, we found no striking main effect of sex at transcriptional level (**Suppl. Table 6)**, at least not beyond X and Y-linked genes that are usually found associated with female and male sampling^20^. We could, however, detect clear differences in phenotype between macrophages from sciatic nerve and DRG. Thus, compared to nerve, MHCII+ myeloid cells from DRG expressed markers commonly associated with resident microglia (Figure 9A and **Suppl. Table 7**).

**Figure 9:**
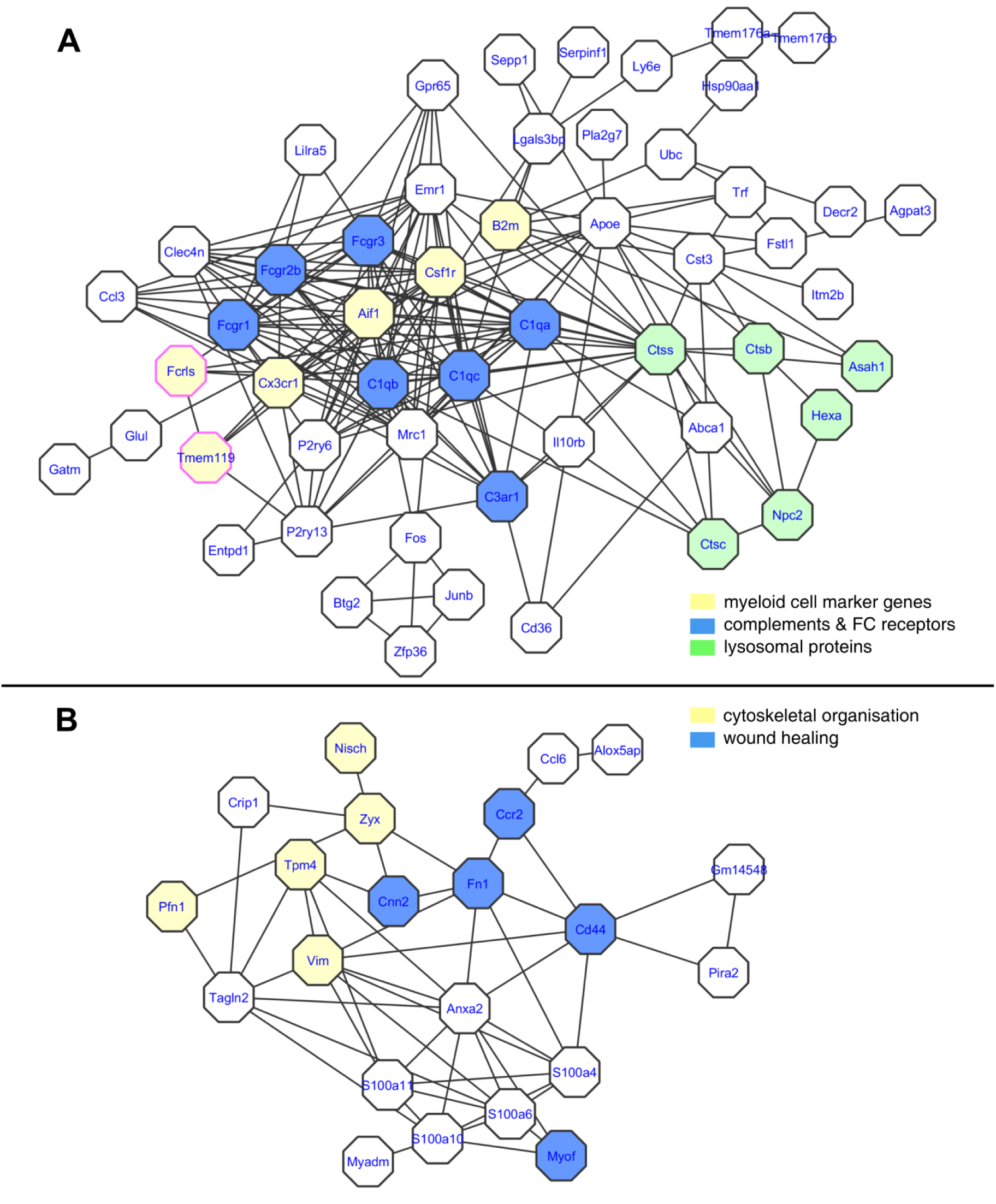
MHCII+/Ly6C-myeloid cells from DRG are different to those from sciatic nerve. **A)** Genes upregulated at adj. p < 0.05 in MHCII+/Ly6C-macrophages from DRG versus nerve were fed into a STRING network analysis. Functional connections were found between 53 (out of a total of 93). Note the presence of canonical microglial marker genes Fcrls and Tmeml 19. **B)** In contrast, genes upregulated in macrophages from nerve rather than DRG formed a very different functional network (22 out of 38) with processes more related to cytoskeletal organisation and wound healing. Note the presence of *Anax2* and several *S100* genes also found at high levels in adipose tissue macrophages (see Figure 11 & Discussion). See sleuth tab in **Supplementary Table** 7 for differential expression.

This microglial-like phenotype is even more striking after injury, in that both MHCII+/Ly6C- and MHCII-/Ly6C-DRG macrophages significantly upregulated a host of known neurodegeneration-associated and homeostatic microglial markers^22^ (Figure 10 and **Suppl. Table 8**). These include *Trem2*, *Apoe, Csf1r, Fcrls, Tmem119* and *Olfml3.* The level of enrichment is highly unlikely to have arisen by chance, with 9x as many neurodegeneration-associated microglial genes identified in DRG as would be expected (p < 0.0001) and 10x as many homeostatic microglial genes (p < 6.5E-12). We validated some of these results with qRT-PCR including additional biological replicates (Suppl. Figure 11). *Trem2* and *Tmem119* were once more significantly dysregulated in DRG compared to nerve in this larger sample, while *Apoe* was not, in line with it being a more borderline result in our sequencing data.

**Figure 10:**
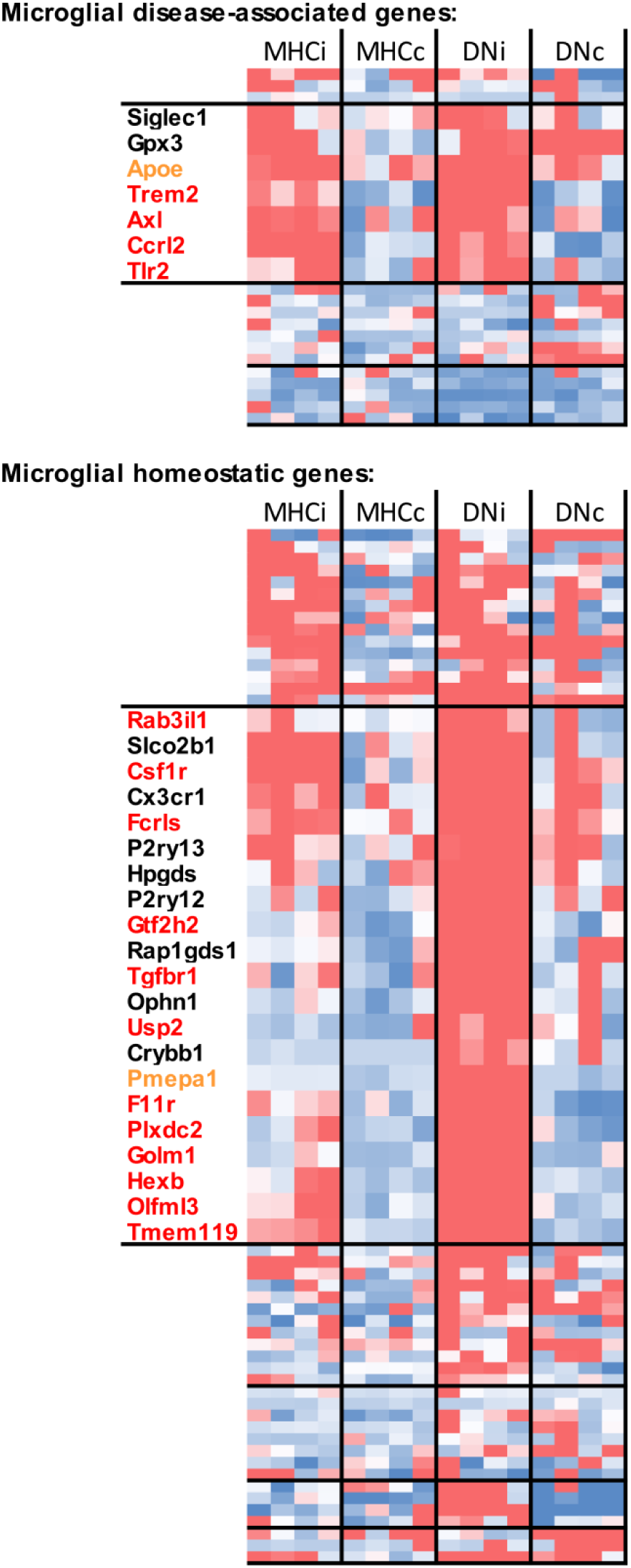
Macrophages in injured dorsal root ganglia up­ regulate genes associated with microglia in homeostasis and disease. Two clustergrams were generated for a list of microglial genes previously shown to be associated with disease vs. homeostasis^22^. This hierarchical clustering of DRG macrophage TPM values revealed blocks of upregulated genes, particularly in the MHCII-/Ly6-resident population. Genes highlighted in red were significantly upregulated in ipsilateral macrophages at q<0.1; genes highlighted in orange at q<0.152. For graphing, values were converted into z space. MHCi: ipsilateral MHCII+/Ly6C-DRG macrophages. MHCc: contralateral MHCII+/Ly6C-DRG macrophages. DNi: ipsilateral MHCII-/Ly6C-DRG macrophages. DNc: contralateral MHCII-/Ly6C-DRG macrophages. See **Supplementary Table 8** for raw data.

With the exception of *Abi3*, *Fscn1*, *Itgax* and *Lgals3*, nerve MHCII+ myeloid cells showed no change in these microglial markers or significantly downregulated them with injury (in the case of *Msr1, Gas7, St3gal6, Scamp5, Csf1r, Cx3cr1, Slco2b1, Gpr34, Siglech*). Compared to DRG, nerve MHCII+ myeloid cells instead expressed relatively more transcripts related to cytoskeletal organisation and wound healing (Figure 9B and **Suppl. Table 7**).

This difference between nerve and ganglion may be a generic one associated with other peripheral neuron-associated macrophages (Figure 11 and **Suppl. Table 7**). A comparison to previously published data on sympathetic neuron-associated macrophages^23^ revealed that genes upregulated in DRG-associated macrophages compared to nerve also appear more highly expressed in macrophages isolated from sympathetic ganglia. The same is true for macrophages from sciatic and sympathetic nerves, both of which resemble adipose tissue macrophages derived from subcutaneous and visceral fat.

**Figure 11:**
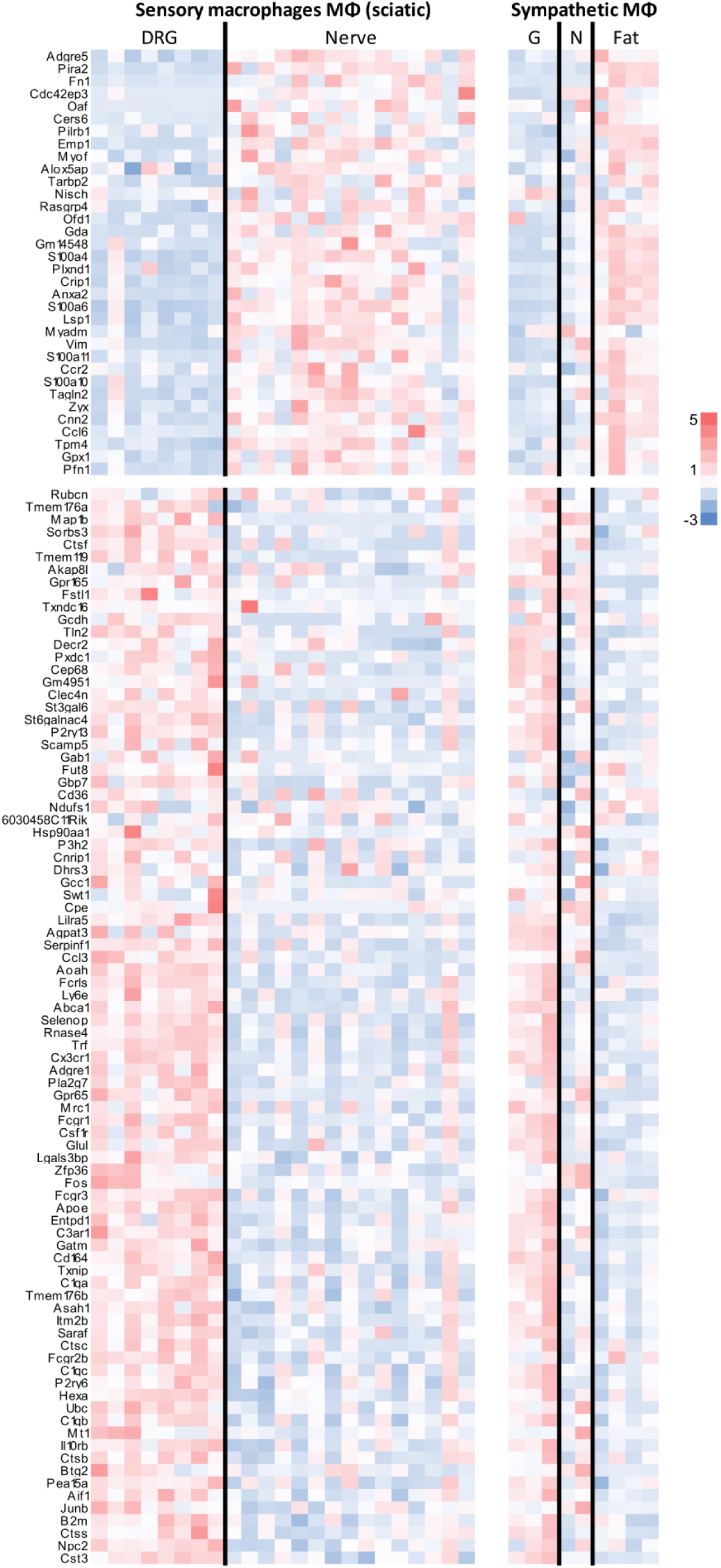
Macrophages from dorsal root ganglia resemble those from sympathetic ganglia, while macrophages from sciatic nerve resemble those derived from sympathetic nerve and adipose tissue. Left: Heatmap of genes significantly up- (top) and downregulated (bottom) in MHCII+/Ly6C-nerve vs. DRG macrophages (q < 0.05). Plotted are TPM values for individual biological replicates, converted into z-scores per gene row. Right: RPKM values for the same list of genes obtained from GSE103847^23^ (again converted into z-scores). MΦ: macrophages. DRG: dorsal root ganglia derived macrophages. Nerve: sciatic nerve derived macrophages. G: macrophages derived from sympathetic ganglia. N: macrophages derived from sympathetic nerve fibres of inguinal fat. Fat: macrophages derived from subcutaneous and visceral fat. See **Supplementary Table 7** for raw data.

**Figure 12:**
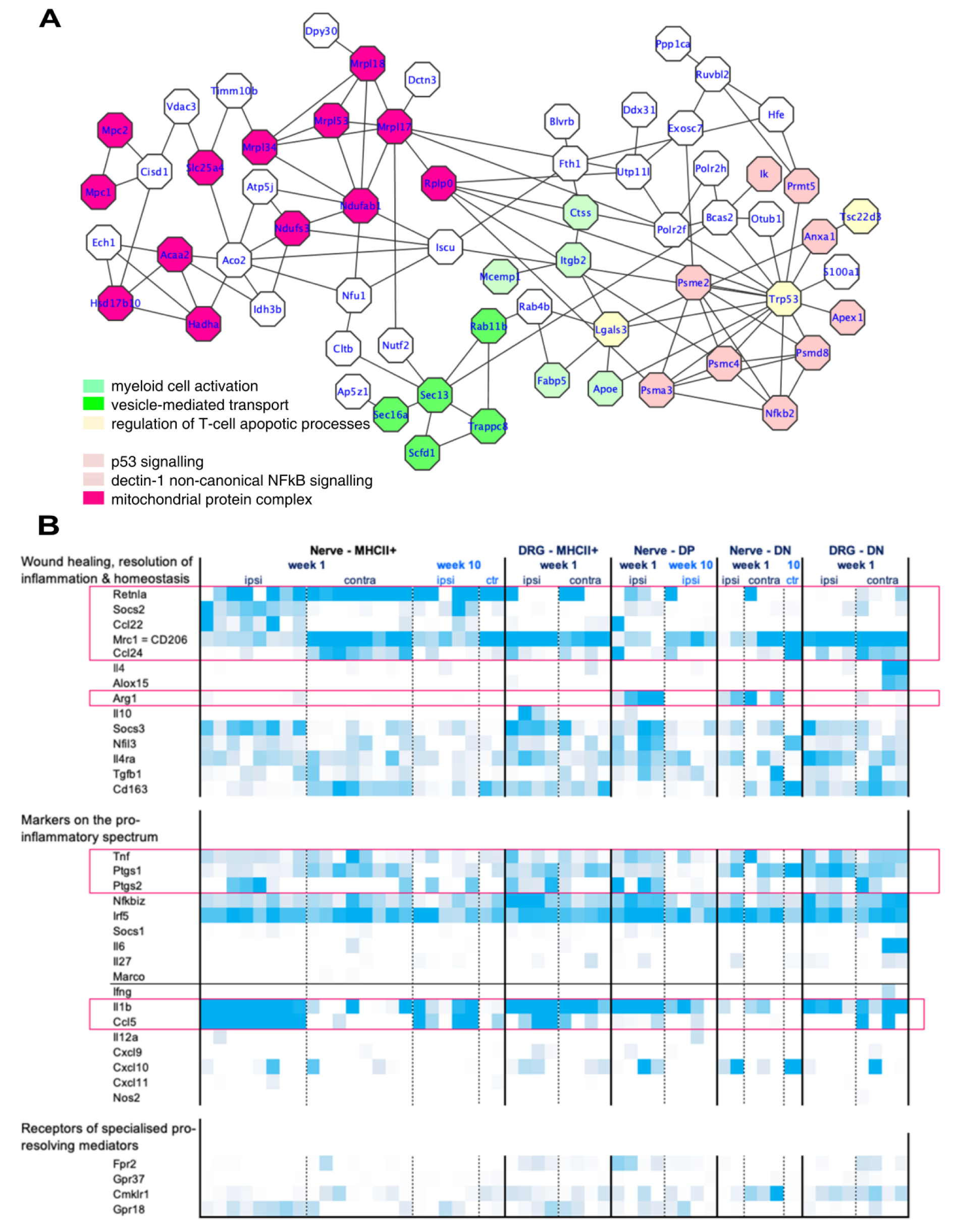
At 2½ months post PSNL, the most prominently upregulated genes in nerve MHCII+/Ly6C-myeloid cells differ from those identified at week 1 - but still do not include transcripts related to resolution of inflammation. **A)** STRING network analysis reveals that 63 of 110 genes upregulated in ipsilateral MHCII+ macrophages at adj. p < 0.05 and FC>3 are likely to be functionally connected with overrepresented processes including myeloid cell activation, vesicle-mediated transport and regulation of T cell apoptosis. See **Suppl. Table 9** for full differential expression tables. **B)** Heat map of genes the expression of which one might expect to see varied with resolution of inflammation. Plotted are TPM expression values from low (white, TPM = 0) to high (darkest blue). MHC+: MHCII+/Ly6C-macrophages. ON: MHCII-/Ly6C-macrophages. DP: MHCII+/Ly6C+ macrophages. See **Suppl. Table 10** for raw data.

10 weeks after PSNL, MHCII+ myeloid cells in sciatic nerve overexpress a different set of genes in response to nerve injury compared to week 1 (Figure 12A and **Suppl. Table 9**). The lists of significantly upregulated genes at the two time points (110 in week 10 vs. 186 in week 1) only have four genes in common – this is not a statistically reliable overlap and could easily arise by chance. Considering the most reliably dysregulated genes at week 10 (FC>3), they appear to comprise transcripts relating to general myeloid cell activation, including *Ctss*, *Lgals3* (galectin-3), *Apoe*, *Fabp5*, *Itgb2* and *Mcemp1*, and genes related to vesicle-mediated transport. A large number of transcripts appear to be related to p53 signalling (*Trp53*) which could point towards cells initiating apoptosis, entering cell cycle or driving chronic inflammatory processes – all functions in which p53 has been implicated^24^. Related to this may be the observed changes in genes involved in the mitochondrial protein complex, some of which control the permeability of mitochondrial membranes during apoptosis (*Slc25a4, Acaa2, Bloc1s2*^1^).

Conspicuously absent in the transcriptional signatures we observed were clear time-dependent changes in genes related to the resolution of inflammation (Figure 12B). Receptors for specialised pro-resolving mediators and known markers for wound healing, resolution of inflammation and macrophage homeostasis were regulated in similar fashion in MHCII+ myeloid cells from ipsilateral nerve – whether they were harvested 1 week or 10 weeks after injury. For instance, *Arg1* and *IL10* were not expressed at either time point, *Mrc1*/CD206 was downregulated at both, and *Socs2* and *Ccl22* upregulated at both. Similarly, pro-inflammatory markers *Il1b* and *Ccl5* remained high over the 2½ months.

With the later phases of inflammation in mind, we also examined the T cell signature at week 10. However, the data provided more questions than answers. There were very few statistically significant changes in T cells from ipsilateral nerve versus contralateral nerve (**Suppl. Table 9**) – possibly because our broad sorting strategy on any βTCR positive cell introduced noise due to T cell subtypes. For instance, it is clearly apparent that one of the ipsilateral samples did contain more CD8a transcript than the others, which were mostly CD4 positive and CD8a negative (Suppl. Figure 9).

It needs to be noted that some of our failure to observe transcriptional changes over time may have been due to lack of sensitivity. Results from the power simulation package *powsimR* indicate that the high dispersion in our week 10 MHCII+ or T cell samples may have meant that we were only powered to detect very large effect sizes (logFC>1.4) at the n numbers we had (Suppl. Figure 12A). In contrast, our MHCII nerve comparisons at week 1 were probably more adequately powered, as our dispersion rates and n numbers should allow for detection of logFC>1 around 80% of the time, depending on expression level (TPM > 10). False positive rates should have been reasonable (below 5%) for all comparisons at all time points for genes expressed at TPM > 10 (Suppl. Figure 12B).

Finally, to integrate the large amount of data we generated with previously published data in an easily accessible fashion, we created a dedicated webpage to browse Neuroimmune Interactions in the Periphery (NIPPY, http://rna-seq-browser.herokuapp.com/). We re-analysed our own data, as well as previously published data on relevant cell types that one might want to consider when examining the expression level of one’s favourite gene in peripheral neuro-immune interactions, particularly after nerve injury. Specifically, we included naïve magnetically sorted sensory neurons (GSE62424)^25^, naïve FAC-sorted sensory neurons (GSE100035)^20^, magnetically sorted sensory neurons after nerve injury (GSE100035)^20^, Schwann cells sorted after sciatic nerve transection (GSE103039)^26^, macrophages after sciatic nerve transection (GSE106488)^27^, CCR2+Cx3cr1+ macrophages after sciatic nerve crush (GSE106927)^28^, and naïve macrophages associated with sympathetic nerve and ganglia in adipose tissue (GSE103847)^23^. To improve inter-study comparisons, we re-aligned our RNA-seq results using BioJupies^29^, to ensure all datasets were processed with the same *kallisto* settings. Our webpage provides a quick overview of expression patterns across murine cell types, as well as a detailed representation of all individual biological replicates within each batch-controlled experiment. Graphs and data can be easily exported. Just to give our readers two example use-cases:

1. a simple search for Il10 reveals that while it is not expressed in our nerve macrophage subsets after partial sciatic nerve ligation (as shown in Figure 12B), it is clearly upregulated in two independent datasets examining its expression level in nerve macrophages after sciatic nerve transection and crush (Figure 13). The latter two are models of nerve regeneration rather than permanent damage and neuropathy. Indeed, IL10 knockout in mice has previously been shown to slow down the speed of resolution and nerve repair after sciatic nerve crush^30^. Moreover, IL10 – if supplemented in models of inflammatory pain – has been shown to be analgesic^31^. Our data indicate that absence of IL10 may be one of the factors that hampers successful resolution of inflammation also in neuropathic pain.
2. looking at expression of the the proenkephalin gene (*Penk*) we noticed that it appears to increase over time in macrophages after sciatic nerve transection (Figure 14A), while it is once more absent in our model of sciatic nerve ligation. We reasoned that maybe *Penk* is specifically upregulated in regenerative states, with macrophages primed to be more wound-healing and less pro-inflammatory. To test this, we treated murine bone-marrow derived macrophages with mediators to mimic either a pro-inflammatory (IFNγ or LPS) or wound-healing (IL-4) activation state. We did this alone or in combination with a neurotransmitter (calcitonin gene-related peptide, CGRP) in order to mimic neuronal activation. *Penk* mRNA was upregulated when the macrophages were treated with IL4 or IL4 and CGRP. This suggests that wound-healing like macrophages, in combination with synergistic signalling from neurons, release proenkephalins (Figure 14B), while macrophages with a more pro-inflammatory-like phenotype do not.

**Figure 13:**
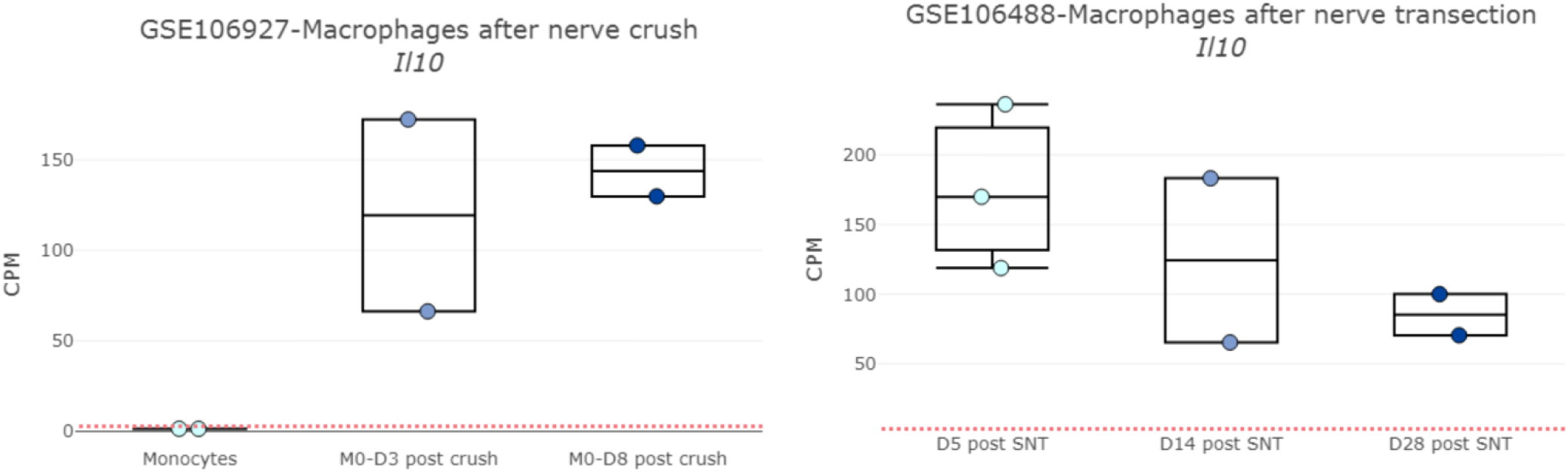
Graphs downloaded with one click from our novel web-resource which allows browsing of cell-type specific RNA-seq data relevant for peripheral neuroimmune interactions.

**Figure 14:**
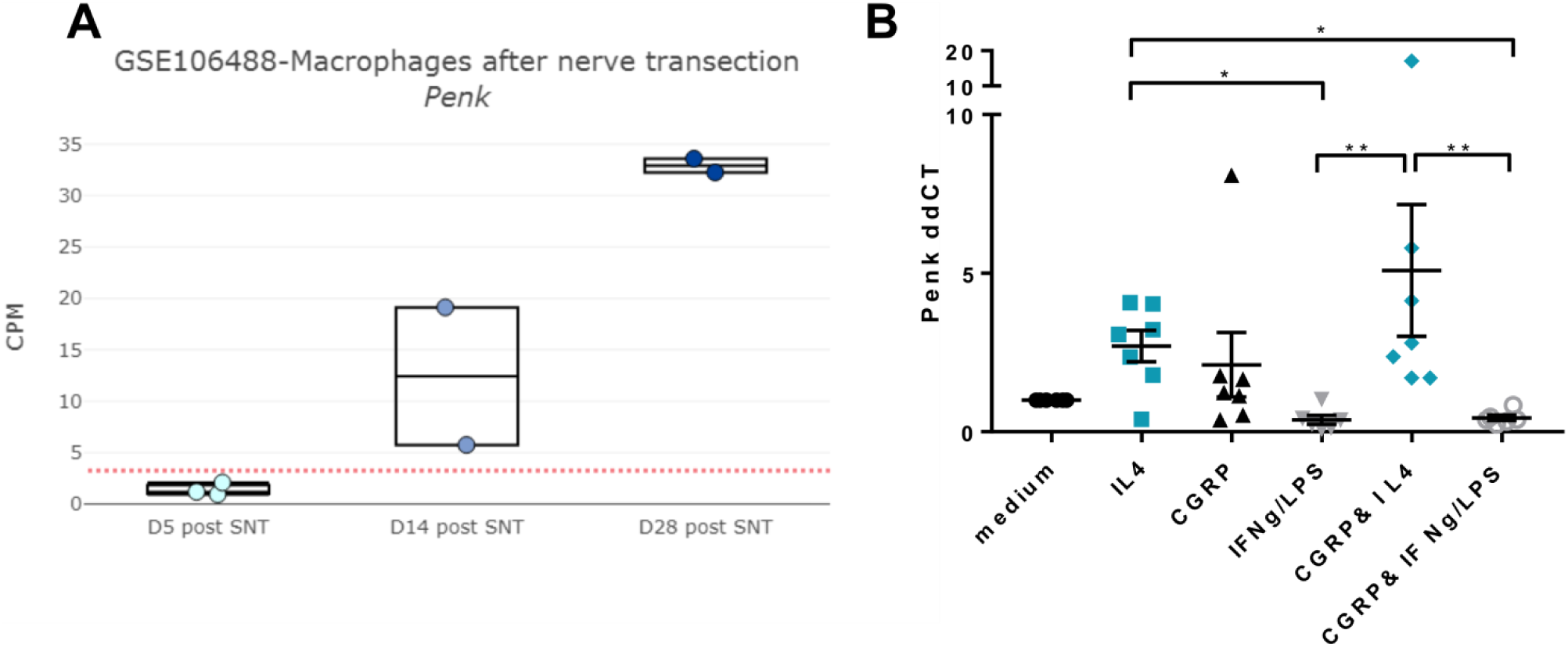
Our web-resource allows formulating and testing of novel hypotheses. **A)** Macrophages isolated from regenerative sciatic nerve after transection appear to progressively increase expression of proenkephalin (*Penk*)- data published under GEO identifier GSE106488 and plotted on our web-resource at counts per million (CPM), i.e. raw counts normalised by the total read count per sample. The dotted red line represents a threshold value below which gene expression is likely to fall within the noise range. Each dot is an independent biological replicate. **B)** qRT-PCR for *Penk* conducted on bone marrow derived macrophages treated with IL4 or the pro-inflammatory cytokines IFNγ or LPS with or without the neuro-transmitter CGRP. IL4 treatment alone and IL4 in combination with CGRP significantly upregulated *Penk* expression: Kursak-Wallis ANOVA X^2^(6) = 25.77, p = 0.0001, with Dunn’s multiple comparisons: *p < 0.03, **p < 0.005. Data points represent values obtained from n = 6-7 independent BMDM culture experiments. Mean ±SEM are also plotted.

## Discussion

Immunologists acknowledge that even “clearly defining inflammation presents a challenge”^18^. Here, we have attempted to move us one step closer toward defining what inflammation means in the context of a model of traumatic nerve injury. Specifically, we reveal the complexity of the immune responses at a cellular and molecular level in both nerve and DRG. We provide evidence to suggest that macrophage sub-populations in nerve and DRG are distinct entities. Sensory nerve macrophages resemble previously published transcriptional signatures of resident macrophages in adipose tissue^23^, while DRG macrophages resemble sympathetic ganglion macrophages^23^ and resident microglia^22^. We demonstrate – in previously unprecedented detail – that resolution of inflammation after nerve injury remains elusive, even after a period of three and a half months. Finally, we identified surprisingly few consistent sex-differences, neither at cellular or transcriptional level.

Our flow cytometry data show that there are two distinct CD11b+/Ly6G-subpopulations in nerve and DRG during homeostasis: MHCII+ single positive and MHCII-/Ly6C-double negative myeloid cells. After nerve injury, these are joined by Ly6C+ cells, likely infiltrating monocytes, and MHCII+/Ly6C+ double positive cells. Our transcriptomic data confirm that MHCII+ single positive cells are antigen-presenting and interacting with lymphocytes. Without lineage tracing it remains unclear whether we are observing antigen-presenting macrophages^32^ or dendritic cells - a subtle distinction that some have anyway claimed to be moot^33^. We can, however, state that the presence of CD45ra receptor transcript (*Ptprc*) together with CD11b, make them unlikely to be blood-derived dendritic cells, at least as defined in a consensus paper on nomenclature in 2010^34^.

Our results further suggest that, at least quantitatively, MHCII+ and MHCII-/Ly6C-macrophages are different in nerve and DRG and react differently after injury. As expected, there is much less proliferation/infiltration of immune cells into DRG than into nerve – the actual site of injury. Moreover, DRG macrophages resemble sympathetic ganglion macrophages and express resident myeloid cell markers, such as *Tmem119* and *Fcrls*, more prominently than nerve macrophages. Conversely, the transcriptome of nerve macrophages is much more similar to that of sympathetic nerve and adipose tissue macrophages. All three cell type, for instance, show relatively more mRNA expression of annexin-2 (*Anxa2*) and its interacting calcium-related adapter proteins (*S100a4, S100a6, S100a10, S100a11*)^35^. Annexin-S100 complexes have been implicated in various cellular functions including endocytosis, cell-cell interactions and regulation of lipids. They have also been shown to be modulated by knockout of the prostaglandin COX-2^36^, the primary target of non-steroidal anti-inflammatory painkillers (NSAIDs). With our current set of data it is impossible to tell whether this similarity is due to nerve and adipose tissue macrophages sharing the same lineage or whether it is merely difficult to isolate macrophages from nerve without contaminating them with those from fat.

After injury, an especially striking difference is the upregulation of neurodegeneration-associated microglial transcripts, namely *Trem2*, *Axl*, *Ccrl2* and *Tlr2*, in MHCII+ DRG, but not nerve macrophages. The same genes are even more starkly dysregulated in MHCII-/Ly6C-populations – very clearly so in DRG, and possibly also in nerve, though the picture is less clear due to the smaller n numbers that were available for sequencing. *Trem2* has been linked to the detection of phosphatidylserine, a membrane-associated lipid that becomes exposed on apopotic or damaged neurons^37^. Evidence suggests that this detection then “tags” the associated cell for microglial phagocytosis^22^. Our results indicate that this is not a microglial-specific process, as is often implied^22, 38^, even in neuroscientific reviews that discuss its presence in macrophages^39^. This is further supported by a very recent report which convincingly argues that high levels of *Trem2* in macrophages is a common feature of pathologies which involve lipid abnormalities, since it has been reported in macrophage sub-populations of atherosclerotic aortas^40^ and of liver and visceral adipose tissue in the context of obesity^41^. An intriguing observation is that Amit et al.^41^ found obesity-associated *Trem2*+ macrophages to derive from blood monocytes, while we find *Trem2* most clearly upregulated in MHCII-/Ly6C-resident subpopulations of the DRG, which, transcriptionally, are least like adipose tissue macrophages and most like resident microglia. Does this mean that resident neuron-associated macrophages have adapted a generic myeloid cell pathway that senses lipid abnormalities to suit their interactions with nerves? Or, as a neuroscientist might be seduced to speculate, is *Trem2*+ macrophage expansion in any tissue to some extent dependent on neuronal abnormalities – which have been reported in the context of both obesity^42^ and atherosclerosis^43^?

Beyond phenotypic identity, the kinetics of the inflammatory response we observed is in keeping with what has previously been reported in the literature. The time-course of neutrophil infiltration into rat nerves was previously characterised using immunohistochemistry^10^. Equivalent to our findings, the authors reported the peak of neutrophil infiltration 24h after PSNL, with fewer, but still noticeably elevated numbers reported 7 days post-PSNL.

Similarly, there have been prior reports of persistent immune cell responses in nerve and DRG after injury, though studies beyond one month are very scarce. To the best of our knowledge, we are only aware of one series of pioneering papers by Hu & McLachlan, who examined macrophage markers in rat nerve and DRG in various models of nerve injury. They reported increased numbers of MHCII+ macrophages and T cells in sciatic nerve and DRG 10 and 11 weeks after L5 spinal nerve transection & ligation^44–46^ – a result which we have now been able to replicate and extend to 14 weeks post sciatic nerve ligation. Impressively, with the help of just three immunohistochemical markers and a careful examination of cell morphology, the group already posited the existence of four distinct macrophage populations in DRG^46^: CD68+/CD163+ resident macrophages, likely equivalent to our MHCII-/Ly6C-population; MHCII+ monocytes invading from blood, likely equivalent to our MHCII+/Ly6C+ population; CD68+/MHCII+ cells, which likely are our MHCII+ single positive population; and CD68+ invading monocytes – equivalent to our Ly6C+ population.

While this information has been available in the literature, and immune cells are generally acknowledged to play a key role in neuropathic pain states^7, 8, 16^, the prevailing narrative in the pain field is still one that segregates pain into inflammatory versus neuropathic^47–51^. It is a clinically useful distinction, since neuropathic pain patients do not respond to non-steroidal anti-inflammatory medications. However, we would argue that, at least from a pre-clinical perspective, this dichotomy evokes dangerously misleading ideas that imply that neuropathic pain states involve “less” inflammation or immune cell activation within the affected tissues. This is emphatically not the case: our data demonstrate that one of the most commonly used animal models of traumatic nerve injury is accompanied by a significant and persistent inflammatory response which is unresolved for 3½ months – for as long as we were permitted to measure it by our ethical review board.

Regarding sex differences, we would like to stress that the absence of consistent sex differences in our flow and RNA-seq data does not mean that they are not present at lower effect sizes that we likely were not powered to detect. We would also like to emphatically endorse the importance of sex and gender considerations when studying neuro-immune interactions – especially since they are known to play prominent roles in the perception of pain, chronic pain conditions and immune system function^52–56^.

Instead, we see our results as a timely reminder of the ease with which a single, conventionally sized experiment (e.g. n = 4 biological replicates) can yield false-positive findings when dealing with variable *in vivo* samples. It is a trap into which we ourselves seem to have fallen^20^. While it can be tricky to estimate power when faced with one-off, small scale, pre-clinical flow-cytometry experiments, it is easier to do so with big RNA sequencing datasets. Our transcriptional experiments were designed to maximise information, by examining a wide variety of cellular phenotypes, and to maximise power, by trading high numbers of biological replicates over depth and single-cell resolution. Nevertheless, with the exception of our MHCII nerve samples at week 1, a combination of shallow read depth and high variance between samples meant that we are only likely to have detected the most striking differences. Moreover, transcripts with TPM < 10 are unlikely to have been adequately represented, both in terms of their expression levels and their variation with injury. It is important to bear this in mind when examining our data – or indeed those of others. Due to cost implications, many published RNA sequencing results will not be adequately powered to detect changes in more lowly expressed genes. Our simulation data indicate that given the high dispersion that is likely be present in the transcriptional data of small numbers of sorted immune cells, you would require at least an n = 16 to be comfortably powered to detect an effect size of logFC>1 at all expression levels (Suppl. Figure 12).

In conclusion, we used a model of traumatic neuropathic pain in order to study peripheral neuro-immune interactions in the context of injury. Our findings point towards a very complex and long-lasting inflammatory response. To help other researchers in this area, we are providing an easy-to-use plug & play website that allows mining of relevant cell-type specific RNA sequencing datasets in naïve and nerve injury conditions.

## Materials and Methods

### Animals

Male and female C57BL/6J mice were purchased from Envigo Ltd at 5-6 weeks of age and acclimatised to the animal unit for one week prior to surgeries. They were housed in standard individually ventilated cages (Tecniplast UK) in groups of five maximum at a 12h light–dark cycle, with ad-lib access to food and water. All experiments described were carried out in accordance with the United Kingdom Home Office Legislation (Scientific Procedures Act, 1986) and were approved by the Home Office to be carried out at King’s College London under project license number P57A189DF.

### Surgeries

Mice underwent partial sciatic nerve ligation surgery, using a paradigm originally developed in the rat^57^. Briefly, animals were anesthetised under isoflurane (Henry Schein, Cat# 988-3245) and administered 0.05 mg/kg of the analgesic buprenorphine subcutaneously. After shaving the fur covering the left hip and leg region, the sciatic nerve was exposed through blunt dissection and roughly half was tightly ligated with a 5.0 suture (Ethicon, Cat# W9982). The wound was stapled shut, and mice were transferred to a clean cage and left to recover in the animal unit. Animals were weighed and monitored regularly (at least tri-weekly) following surgery.

### Processing of sciatic nerve and DRG

Mice were administered an intraperitoneal overdose of pentobarbital (Euthatal; Merial, Lot# P02601A) and perfused with 10 ml of 1 x PBS to avoid peripheral blood contamination. The ligated sciatic nerve was exposed (Suppl. Figure 1) and followed up towards the spinal cord to identify and dissect out L3-L5 DRG into F12 (Gibco, Cat# 21765-029). The nerve itself was also placed into a petri dish containing F12 and cut to 0.5 cm, ensuring equal lengths of nerve either side of the ligature. If still present, the ligature itself was removed with tweezers. This process was repeated for the uninjured contralateral side, with pains being taken to ensure harvesting from a similar segment of nerve (again 0.5 cm in length). Tissues were kept in F12 on ice until all animals in a given batch were processed and then transferred into wells of a standard PCR plate (Starlab, Cat# E1403-0100) containing 50 μl of digestion mix: F12 with 6.25 mg/ml collagenase type IA (Sigma Aldrich, Cat# C9891), 0.4% w/v hyaluronidase (ABNOVA, Cat# P52330) and 0.2% w/v pronase (Millipore Cat# 53702) for nerve; F12 with 3mg/ml dispase ll (Sigma Aldrich, Cat# 04942078001), 12.5mg/ml collagenase type IA (Sigma Aldrich, Cat# C9891) and 10mg/ml DNase l (Sigma Aldrich, Cat# 10104159001) for DRG. For optimal digestion, nerves were chopped into small pieces with spring scissors (50x chops, cleaning scissors with 100% ethanol between samples) and incubated at 37°C, shaking at 220 RPM for 30 mins. The digestion mix was subsequently removed and replaced with 100 μl of FACS buffer: HBSS (Gibco, Cat# 14175095) with 0.4% bovine serum albumin (Sigma-Aldrich Cat# A3983), 15 mM HEPES (Gibco Cat# 15630080) and 2 mM EDTA (Invitrogen Cat# 15575-038). Samples were dissociated by pipetting up and down 50x and passed through 70 μm Flowmi cell strainers (SP Scienceware, Cat# 136800070) into a 96 well v-bottom plate. The plate was centrifuged at 1500 RPM for 5 minutes at 4°C, supernatant was removed, and the pellets then underwent antibody staining as described in the following.

### Flow Cytometry/ FACS

On ice, cells were incubated in fixable viability dye in HBSS for 15 minutes, followed by 15 minute incubation in a mix of directly conjugated antibodies and Fc block (see Suppl. Table 1 for antibody panels and concentrations). After staining, cells were centrifuged for 5 minutes (1500 RPM at 4°C) and supernatant was removed. For flow cytometry experiments only, pellets were fixed in 4% PFA for 5 minutes, the plate was then centrifuged for 5 minutes (1500 RPM at 4°C) and supernatant was discarded. Finally, pellets were re-suspended in 200 μl of FACS buffer. Flow cytometry and FACS were performed at the NIHR BRC flow core facility at King’s College London on a BD Fortessa and BD FACSAria, respectively. Unstained cells and single staining control beads (BD Bioscience, Cat# 552845 for antibodies; Invitrogen, Cat# A10346 for the viability dye) were used for compensation. Fluorescence-minus one controls were recorded and used for gating (see Suppl. Figures 5 & 6 for gating strategies employed). For FACS experiments 20 cells were collected per population of interest. Analyses were carried out using FlowJo software.

### RNA sequencing

RNA from FACS collected samples was reverse transcribed into cDNA, amplified and purified in accordance with the Smart-seq2 method^58^. Samples were then run on a High Sensitivity DNA chip (Agilent, Cat# 5067-4626) on a 2100 Bioanalyzer (Agilent, Cat# G2939BA) to determine quality and concentration. SMARTer amplified cDNA was shipped to the Oxford Genomics Centre for batch-controlled library preparation using an Illumina NexteraXT kit (Illumina, Cat# FC-131-1096) and sequenced on an Illumina HiSeq 4000 (75 bp, paired-end reads).

### Data analysis

Basic quality control was conducted in seqmonk on stampy bam files^59^ obtained from the sequencing facility. For detailed analysis, fastq files were trimmed by Trimmomatic^60^ (version-0.39) to eliminate reads with phred scores less than 33 and remove any residual adapters. The reads were then pseudo-aligned using kallisto^61^ (version 0.46.0) with the following command parameters: *kallisto quant-i transcripts.idx -o output -b 100* [*lane1*]*.fastq* [*lane2*]*.fastq* [*lane3*]*.fastq.* The *transcripts.idx* file was built from Ensembl v96 transcriptomes according to instructions provided by the Pachter lab (https://github.com/pachterlab/kallisto-transcriptome-indices/releases). Raw transcript counts were length and depth corrected by conversion to transcripts per million (TPM) using the R package tximport^62^ and the transcripts_to_genes file provided on the kallisto website. The R package sleuth^21^ was used for differential expression analysis (Wald test for two-group comparisons, likelihood ratio test for multiple group comparisons). Power simulations were conducted using the R package powsimR^63^. Data were deposited in the Gene Expression Omnibus (GEO) archive, accession number GSE139150.

To generate our web-tool, we also ran our data through kallisto on BioJupies^29^ to be able to compare our results to those of other published cell-type specific datasets in the same format. BioJupies is a re-analysis tool based on ARCHS4^64^ – a project with one common kallisto pipeline to re-align all human and mouse RNA-seq experiments which were run on HiSeq 2000, HiSeq 2500 and NextSeq 500 platforms and deposited on either GEO or the Short Read Archive (SRA). We accessed cell types analysed in their naïve state, in regeneration models (sciatic nerve transection and crush) and in persistent pain models (sciatic nerve ligation). In particular, we compiled: naïve magnetically sorted sensory neurons (GSE62424)^25^, naïve FACS-orted sensory neurons (GSE100035)^20^, Schwann cells sorted after sciatic nerve transection (GSE103039)^26^, macrophages after sciatic nerve transection (GSE106488)^27^, CCR2+Cx3cr1+ macrophages after sciatic nerve crush (GSE106927)^28^, naïve macrophages associated with sympathetic nerve and ganglia in adipose tissue (GSE103847)^23^.

### Behavioural assay

Behavioural data was collected at ∼13.5 weeks following PSNL. To avoid bias, behaviours were video-taped and later scored by two individuals unfamiliar with pain models and our hypothesis, and who were unable to distinguish between male and female mice. For the test, mice were left to freely explore an open arena of 20cm in diameter – once for five minutes two days prior to testing without video recording for acclimatisation and once for two minutes with video recording. Videos were scored for deficits to the ipsilateral and contralateral legs and hind-paws using a simple 0-2 scale, whereby 0 = no deficit, 1 = slight paw clenching and 2 = severe paw clenching and/or dragging of leg. Numbers were awarded every 30 seconds so that each mouse received a total deficit score out of 8. Scoring was performed on each hind leg separately. Final scores were aggregated via averaging of the individual scores obtained by the two independent observers.

### Immunohistochemistry

At 14 weeks post PSNL surgery, mice were overdosed with pentobarbital and perfused with 10 ml of 1 x PBS, followed by 20 ml of 4% paraformaldehyde (PFA). 0.5 cm of ipsilateral and contralateral sciatic nerves were dissected as described above for flow cytometry. Nerves were then post-fixed in 4% PFA for 2 hours, washed in 1 x PBS and cryoprotected in 30% sucrose. Following this, nerves were embedded in OCT (CellPath, KMA-0100-00A) and cut sagittally into 20 µm sections on a cyrostat. For immunofluorescent staining, sections were incubated for 4 hours in blocking serum (10% donkey serum, 0.2% triton in PBS) and stained overnight with primary antibodies against IBA1 (rabbit, 1:500, Wako, 019-19741), CD68 (mouse, 1:500, Abcam, ab31630) and directly conjugated MHCII (1:500, BioLegend, 107607). After 3 washes in PBS, sections were incubated with secondary antibodies for 2 hours – AlexaFluor647 (goat anti-rabbit, 1:1000, Invitrogen, A21244) and AlexaFluor568 (donkey anti-mouse, 1:1000, Invitrogen, A10037). After 3 further washes in PBS, sections were mounted using Fluoromount-G mounting medium with DAPI (Invitrogen, 00-4959-52) and imaged on a Zeiss LSM 710 confocal microscope.

### BMDM culture & qRT-PCR

Bone marrow-derived macrophages (BMDMs) were isolated from C57BL/6J mice and cultured in Petri dishes containing Dulbecco’s Modified Eagle’s Medium (DMEM) (Gibco, 32430-027) supplemented with 10% (v/v) fetal bovine serum (10500-064) and Pen/Strep (Sigma Aldrich, P4333). Femur and tibia bones from both hind limbs of female mice were collected and the bone marrow was flushed out with cold PBS (Gibco, 14190-094) under sterile conditions using a 10ml syringe with a 25G needle. Cells were centrifuged (7 min, 2000 rpm, room temperature) and the pellet was resuspended in DMEM-MCSF (Peprotech, 315-02). The suspension was passed through a 40 μm cell strainer to filter out bone fragments. Flow-through was collected and filled up to 50 ml with pre-warmed DMEM-MCSF. The cell suspension was plated onto two 15 cm Petri dishes (25 ml/dish) and incubated (37 °C, 5% CO2) for 7 days to allow for differentiation into mature naïve macrophages. After 7 days of incubation, the culture supernatants were discarded and the remaining adherent cells were washed with 15 ml of pre-warmed PBS. Cells were incubated (37 °C, 5% CO2) with 15 ml of pre-warmed non-enzymatic cell dissociation solution (Specialty Media, CAT#S-014-B) for 10 min and then gently dislodged with a cell scraper and centrifuged for collection (7 min, 1800 rpm, room temperature). The cells were plated at 5×10^5/well in 12 multi-well plates or 24 multi-well plates and left to settle overnight in DMEM (w/o MCSF). The next day, cells were incubated for 24 hours in: IFNγ 50 ng/ml (Peprotech, 315-05) or LPS 100 ng/ml (Sigma, L4391) or IL4 50 ng/ml (Milteny biotech, 130-097-760), either alone or in combination with CGRP 100 ng/ml (Bachem, H-1470). Unstimulated cells were incubated in plain DMEM as a control. For RNA extraction, BMDMs were lysed (RLT-buffer+ b-mercapto-ethanol) and RNA was extracted using the RNAeasy Microkit 50 (Qiagen, 74004) according to manufacturer’s instructions. mRNA quantity was evaluated using Qubit 3 Fluorometer (Invitrogen) or Bioanalyzer (Agilent RNA pico chip). 0.124 – 1.548 ug of RNA was converted into cDNA using SuperScript III Reverse Transcriptase (Invitrogen, 18080-044) according to manufacturer’s instructions. 6.2ng/ul of cDNA was used in standard SYBR Green (Roche, 39355320) qRT-PCR reactions on a LightCycler 480 (Roche) to quantify expression of Penk (forward GAGAGCACCAACAATGACGAA; reverse TCTTCTGGTAGTCCATCCACC). The geometric mean of the housekeeping gene 18s was used to calculate the 2^-ΔCt values plotted in Figure 14B.

### qRT-PCR validation of sequencing data

The following primer sequences were used to assess the expression level of Apoe, Trem2 and Tmem119 in additional biological replicates of MHCII+ nerve/DRG macrophages and MHCII-/Ly6C-DRG macrophages. All primers, including the Penk primer above, were tested for their efficiency and specificity:

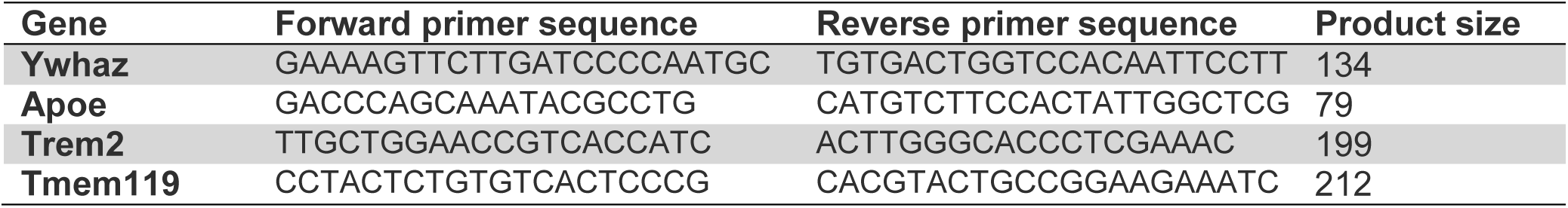

## Acknowledgements

This work has received funding from an MRC New Investigator Grant (MR/P010814/01) and the Innovative Medicines Initiative 2 Joint Undertaking under Grant Agreement n° 116072. This Joint Undertaking receives the support from the European Union’s Horizon 2020 research and innovation. We thank the Oxford Genomics Centre at the Wellcome Centre for Human Genetics (funded by Wellcome Trust grant reference 203141/Z/16/Z) for the generation and initial processing of the sequencing data. The authors acknowledge financial support from the Department of Health via the National Institute for Health Research (NIHR) comprehensive Biomedical Research Centre award to Guýs & St ThomaśNHS Foundation Trust in partnership with Kinǵs College London and Kinǵs College Hospital NHS Foundation Trust.

---

**Supplementary Figure 1:**
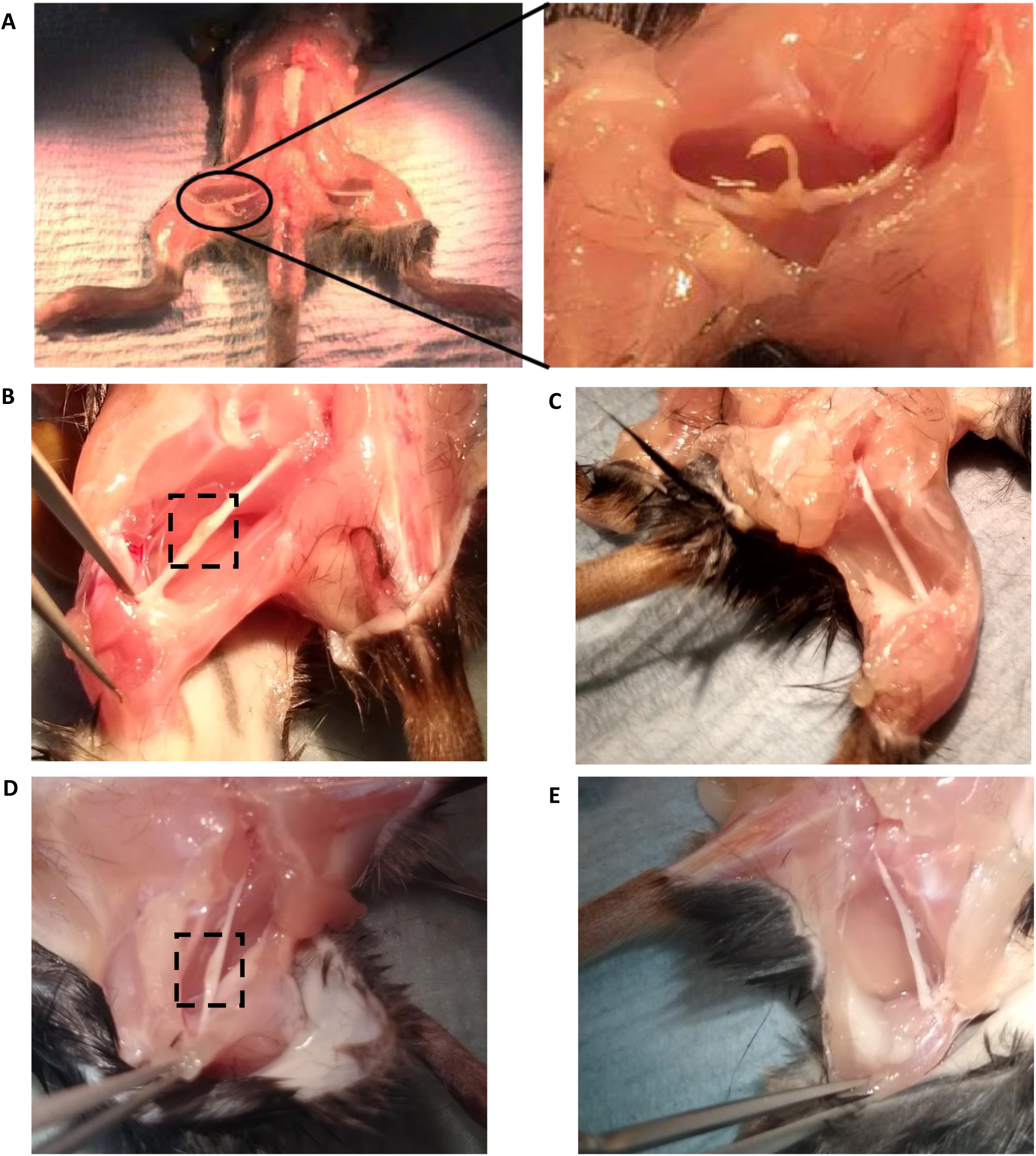
2-3 months after partial sciatic nerve ligation, the suture has been completely absorbed. Representative images of a mouse sciatic nerve after PSNL. **A)** Seven days after surgery, the suture placed through the nerve is clearly visible. **B)** 10 weeks after surgery only a small area of swelling of yellowish hue remains visible on the ipsilateral side. The uninjured contralateral side is shown for comparison **(C)**. **D)** Slight swelling and discoloration is still visible 14 weeks after surgery compared to the contralateral side **(E)**.

**Supplementary Figure 2:**
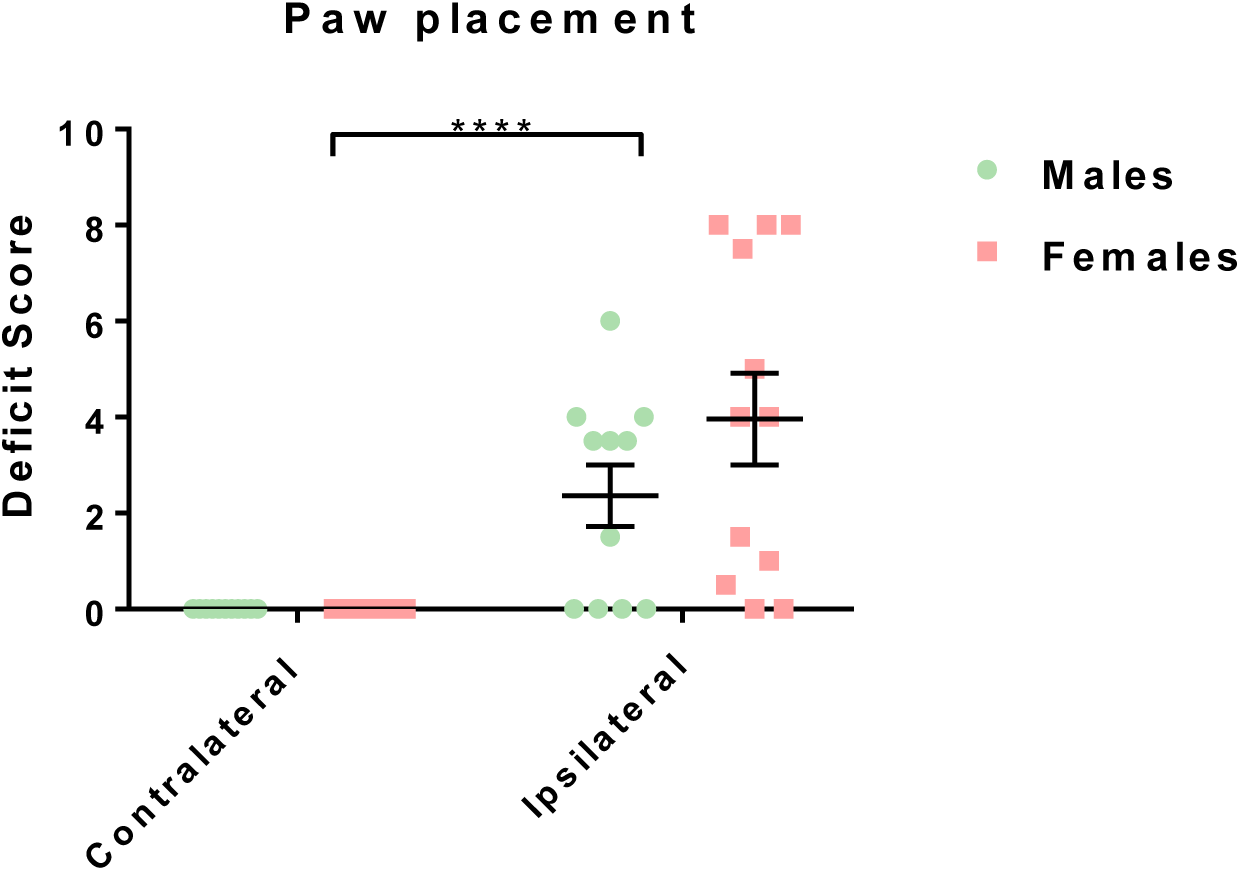
14 weeks post PSNL mice still display behaviours indicating pain in their ipsilateral paw. Two independent observers unfamiliar with pain models, the specific experimental question and its expected outcomes scored videotaped paw placement of mice in an open field over a period of 2 minutes. A two-way repeated measures ANOVA revealed a main effect of paw (ipsi-vs. contralateral): F(1,21) = 28.97, p < 0.0001, which was significant for both male (Sidak post-hoc test, p = 0.02) and female (p = 0.0002) mice. n = 11 male and n = 12 female mice were tested, with scores for each shown here as separate data points with mean and SEM.

**Supplementary Figure 3:**
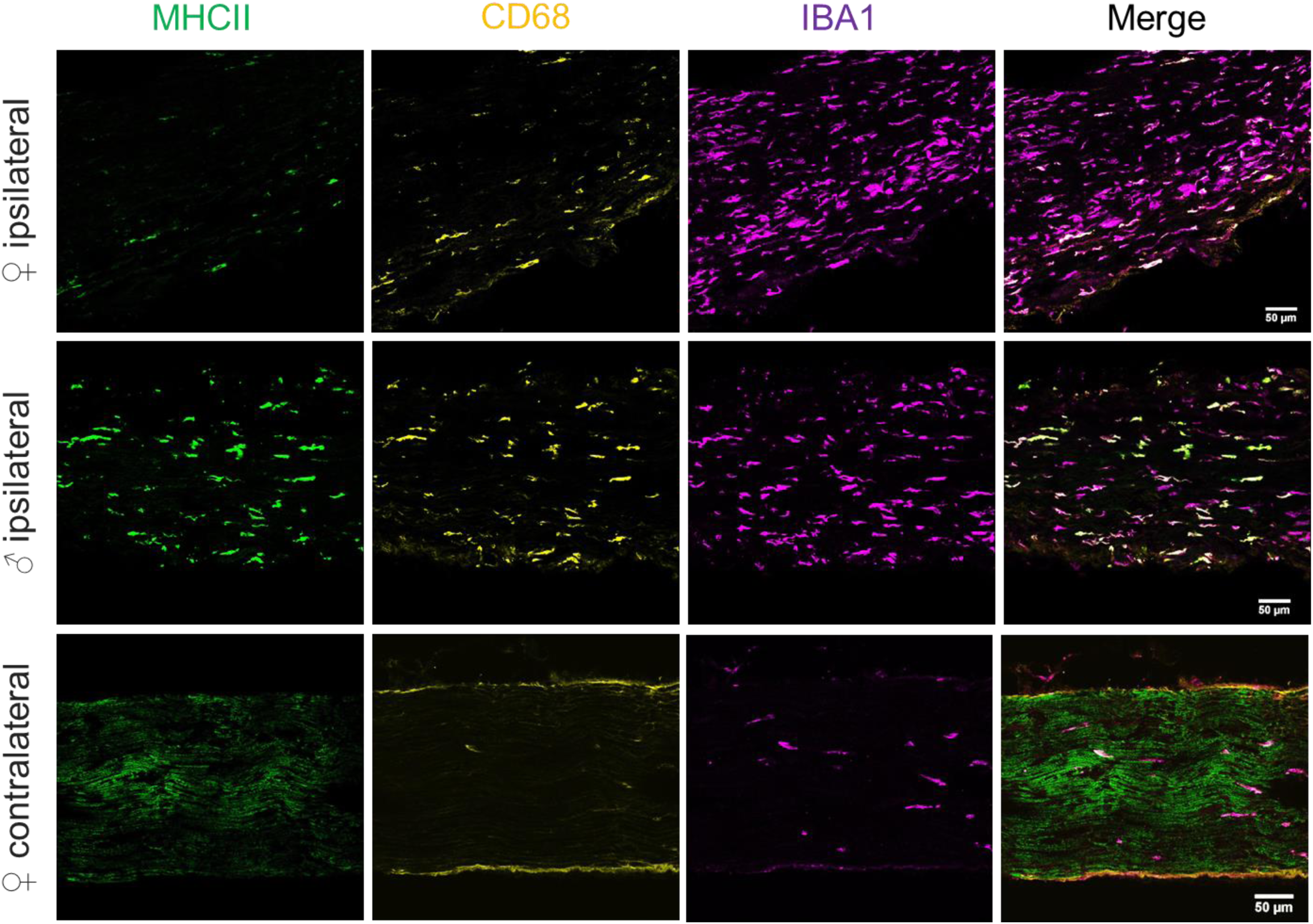
Mirroring our flow cytometry data, immunostaining likewise shows increased expression of macrophages and MHCII+ antigen presenting cells at 14 weeks following injury. Representative images of ipsilateral and contralateral sciatic nerve cryosections from C57B/6J mice of both sexes at 14 weeks post-PSNL surgery. Sections were stained with antibodies against MHCII (green), CD68 (yellow) and IBA1 (purple). Scale bar = 50 μm.

**Supplementary Figure 4:**
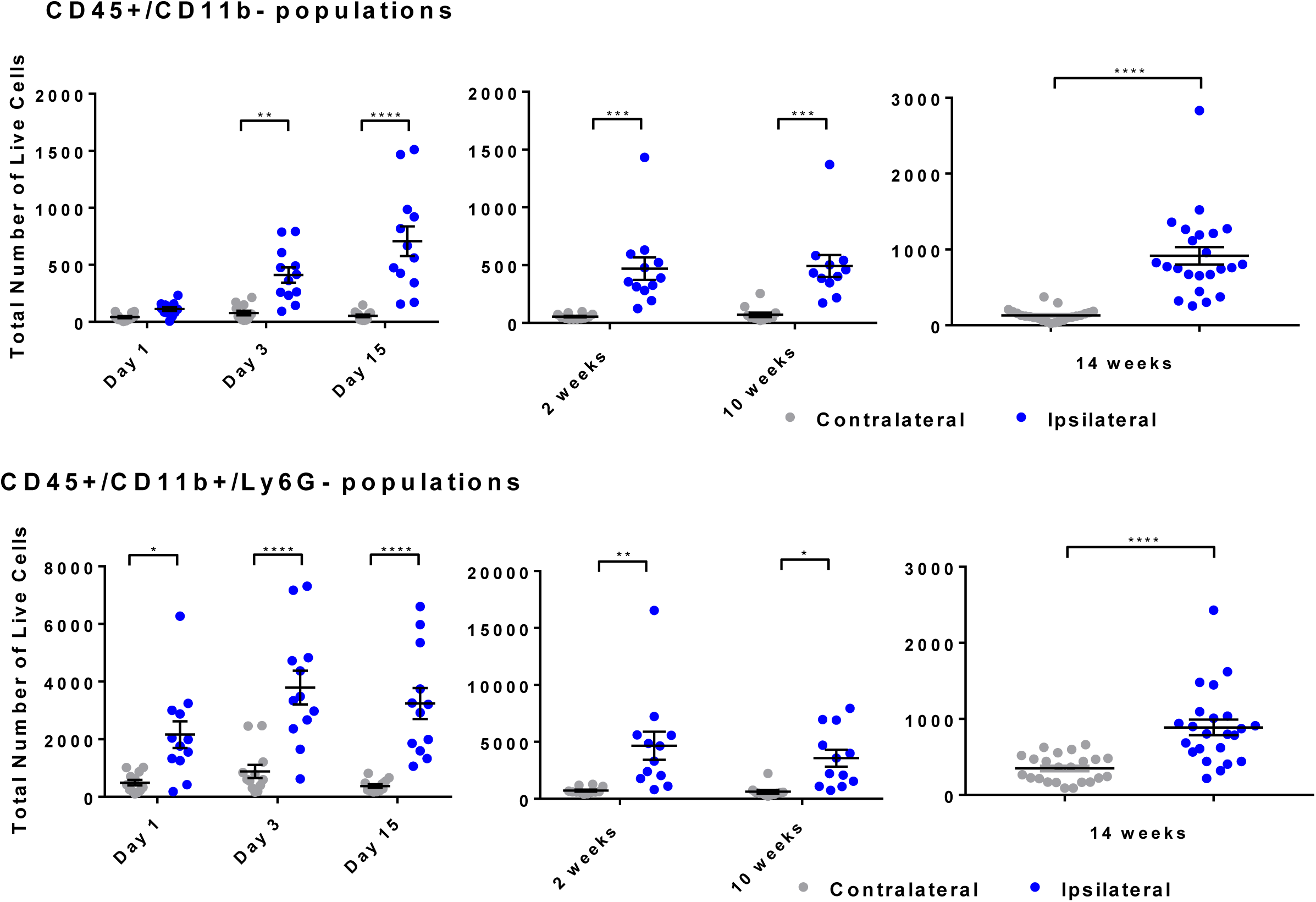
Both myeloid and non-myeloid populations are persistently increased in ipsilateral sciatic nerve until at least 14 weeks post PSNL. **CD45+/CD11b-populations** (lymphoid and other non-myeloid) are significantly upregulated from day 3 post PSNL-onwards, and remain upregulated at comparable numbers 2 weeks and 10 weeks after injury. 14 weeks after injury, the absolute number of events recorded was even higher (∼1000 vs. 500 at 2 & 10 weeks), but given that these experiments were not conducted in the same batch, it is unclear whether this constitutes a biologically significant upregulation over time. **CD45+/CD11b+/Ly6G-myeloid cell populations** show a very similar pattern, with significant upregulation from day 1 PSNL which lasts up to 3 and a half months later. The size of the increase is comparable 2 weeks and 10 weeks after injury, but appears lower 14 weeks after injury in absolute terms. Again, we cannot comment on whether this equates to reduction in inflammation, or whether this is due to differences in experimental batches. At each time point, n = 12 mice were processed (6 males & 6 females), shown here as separate data points with mean and SEM.

**Supplementary Figure 5:**
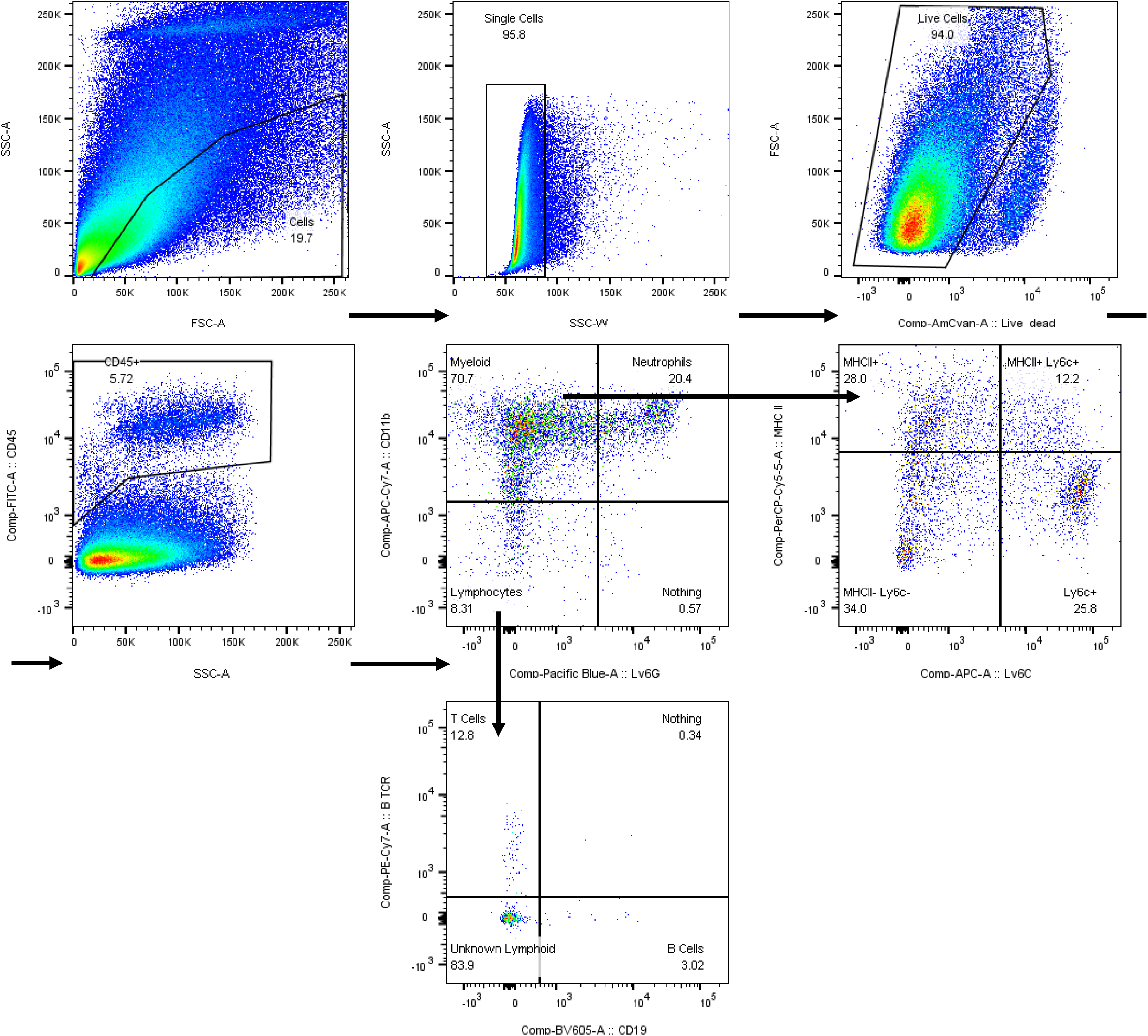
Gating strategy used for first batch of flow cytometry experiments (time points day 1 – day 15). Single live CD45 positive cells were further split into CD11b+/Ly6G-, CD11b+/Ly6G+ and CD11b-/Ly6G-populations. CD11b+/Ly6G-were further split into four populations: MHCII+ single positive (putative antigen presenting macrophages), MHCII-/LyC6-double negative (putative resident macrophages), Ly6C+ single positive cells (likely infiltrating monocytes), and MHCII+/Ly6C+ double positive cells (only present during injury). CD11b-/Ly6G-populations were further split into βTCR expressing T cells and CD19-expressing B cells.

**Supplementary Figure 6:**
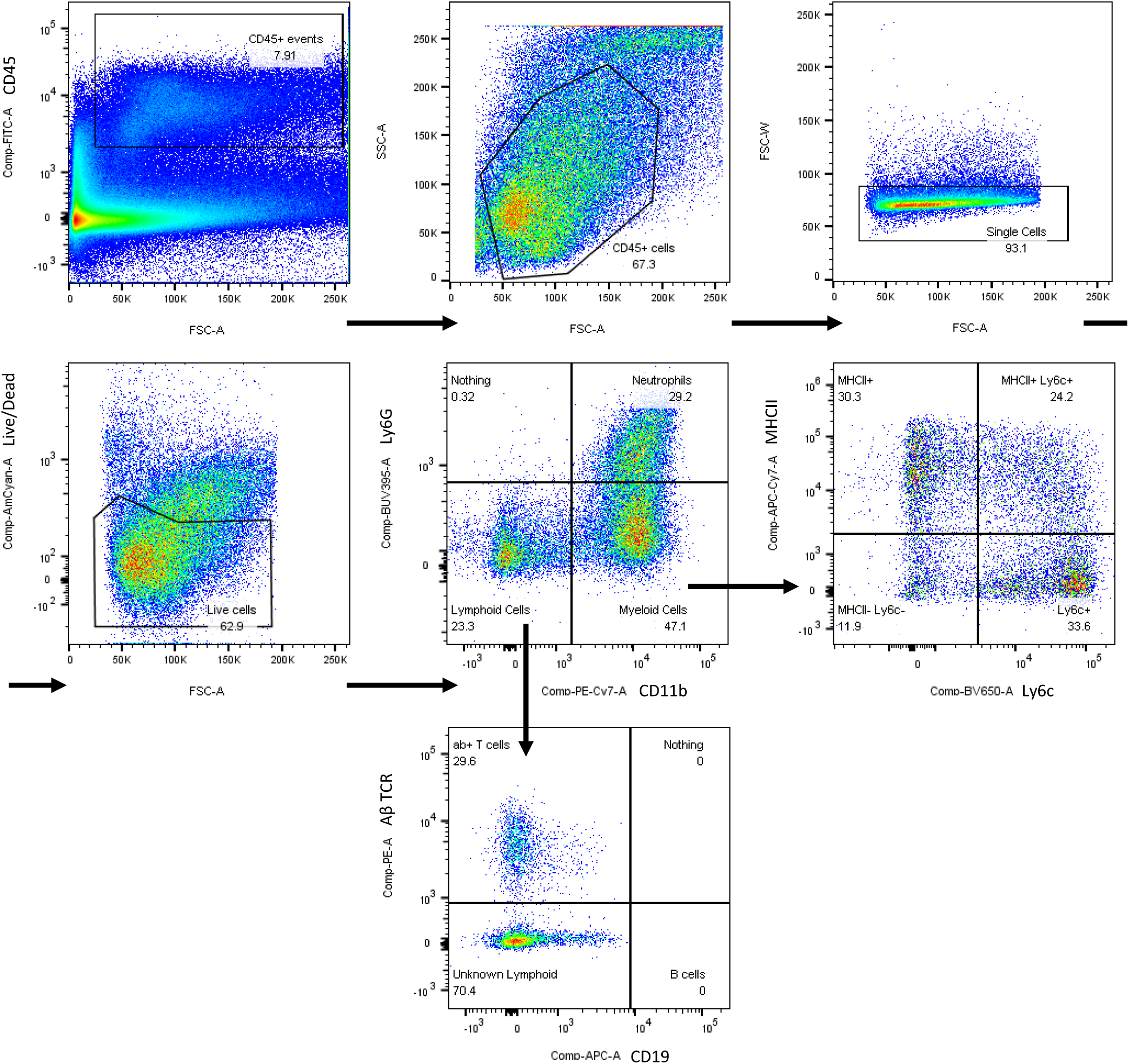
Gating strategy used for all other flow and FACS experiments. Live single cells were selected from CD45 positive cells, which were then further split into CD11b+/Ly6G-, CD11b+/Ly6G+ and CD11b-/Ly6G-populations. CD11b+/Ly6G-were further split into four populations: MHCII+ single positive (putative antigen presenting macrophages), MHCII-/LyC6-double negative (putative resident macrophages), Ly6C+ single positive cells (likely infiltrating monocytes), and MHCII+/Ly6C+ double positive cells (only present during injury). CD11b-/Ly6G-populations were further split into αβTCR expressing T cells and CD19-expressing B cells. *NB: βTCR and CD19 were omitted from the panel in the 2 vs. 10 week flow and 1 week FACS experiments*.

**Supplementary Table 1:**
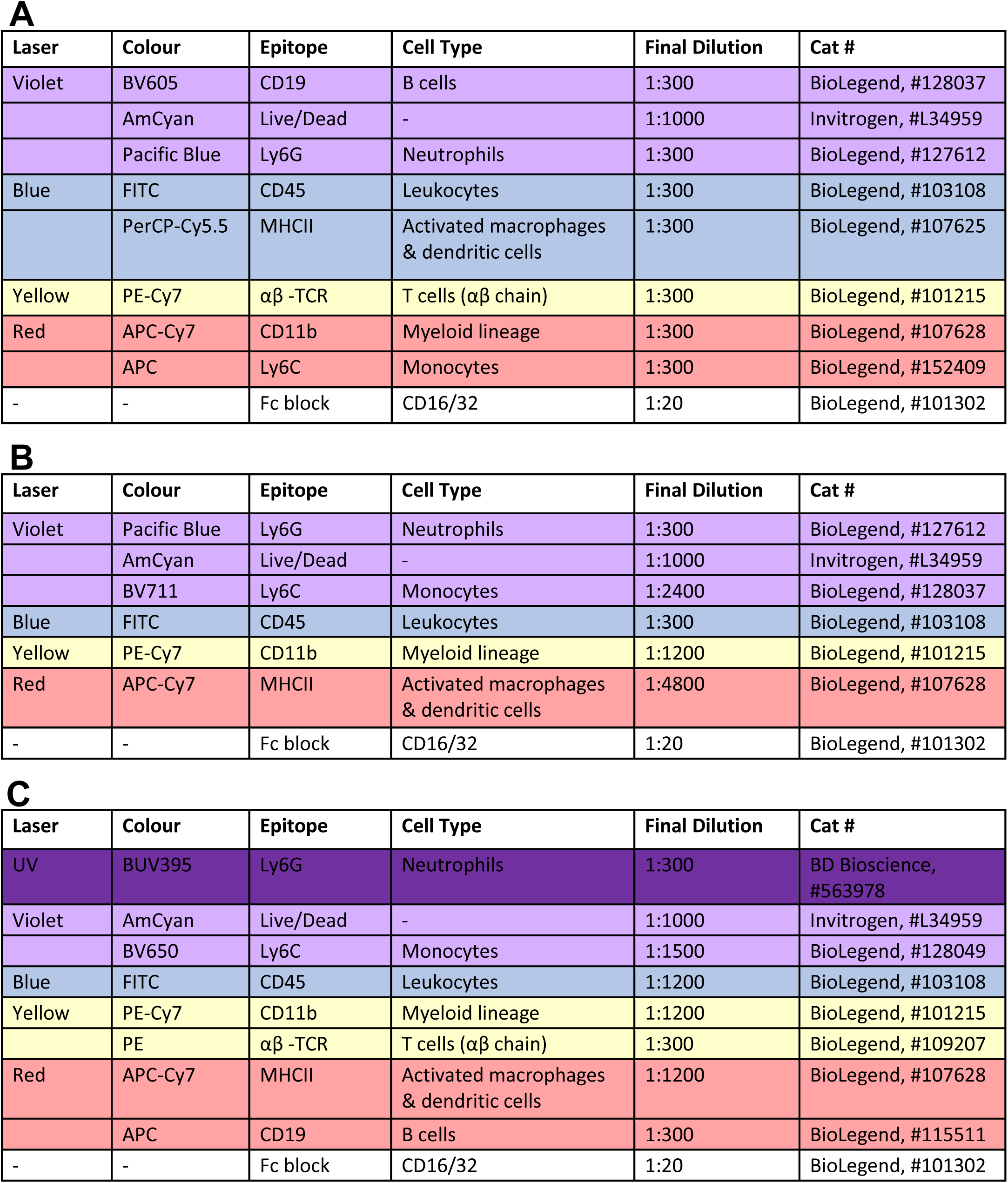
Panels used for flow cytometry and FACS experiments. Experiments were conducted over the span of a year and a half, over which our panels evolved. **A)** Original panel used for 1 day to 2 weeks post-PSNL flow cytometry **B)** Updated panel used for 1 week post-PSNL FACS and 2 vs. 10 weeks post-PSNL flow cytometry experiments. **C)** Final panel – our current choice - used for 10 weeks post-PSNL FACS and 14 weeks post-PSNL flow cytometry experiments.

| Laser | Colour | Epitope | Cell Type | Final Dilution | Cat # |
| --- | --- | --- | --- | --- | --- |
| Violet | BV605 | CD19 | B cells | 1:300 | BioLegend, #128037 |
|  | AmCyan | Live/Dead | - | 1:1000 | Invitrogen, #L34959 |
|  | Pacific Blue | Ly6G | Neutrophils | 1:300 | BioLegend, #127612 |
| Blue | FITC | CD45 | Leukocytes | 1:300 | BioLegend, #103108 |
|  | PerCP-Cy5.5 | MHCII | Activated macrophages & dendritic cells | 1:300 | BioLegend, #107625 |
| Yellow | PE-Cy7 | $\alpha\beta$ -TCR | T cells ($\alpha\beta$ chain) | 1:300 | BioLegend, #101215 |
| Red | APC-Cy7 | CD11b | Myeloid lineage | 1:300 | BioLegend, #107628 |
|  | APC | Ly6C | Monocytes | 1:300 | BioLegend, #152409 |
| - | - | Fc block | CD16/32 | 1:20 | BioLegend, #101302 |

**B**
| Laser | Colour | Epitope | Cell Type | Final Dilution | Cat # |
| --- | --- | --- | --- | --- | --- |
| Violet | Pacific Blue | Ly6G | Neutrophils | 1:300 | BioLegend, #127612 |
|  | AmCyan | Live/Dead | - | 1:1000 | Invitrogen, #L34959 |
|  | BV711 | Ly6C | Monocytes | 1:2400 | BioLegend, #128037 |
| Blue | FITC | CD45 | Leukocytes | 1:300 | BioLegend, #103108 |
| Yellow | PE-Cy7 | CD11b | Myeloid lineage | 1:1200 | BioLegend, #101215 |
| Red | APC-Cy7 | MHCII | Activated macrophages & dendritic cells | 1:4800 | BioLegend, #107628 |
| - | - | Fc block | CD16/32 | 1:20 | BioLegend, #101302 |

**Supplementary Table 1: Panels used for flow cytometry and FACS experiments**
| Laser | Colour | Epitope | Cell Type | Final Dilution | Cat # |
| --- | --- | --- | --- | --- | --- |
| UV | BUV395 | Ly6G | Neutrophils | 1:300 | BD Bioscience, #563978 |
| Violet | AmCyan | Live/Dead | - | 1:1000 | Invitrogen, #L34959 |
|  | BV650 | Ly6C | Monocytes | 1:1500 | BioLegend, #128049 |
| Blue | FITC | CD45 | Leukocytes | 1:1200 | BioLegend, #103108 |
| Yellow | PE-Cy7 | CD11b | Myeloid lineage | 1:1200 | BioLegend, #101215 |
| | PE | $\alpha\beta$ -TCR | T cells ($\alpha\beta$ chain) | 1:300 | BioLegend, #109207 |
| Red | APC-Cy7 | MHCII | Activated macrophages & dendritic cells | 1:1200 | BioLegend, #107628 |
|  | APC | CD19 | B cells | 1:300 | BioLegend, #115511 |
| - | - | Fc block | CD16/32 | 1:20 | BioLegend, #101302 |

**Supplementary Figure 7:**
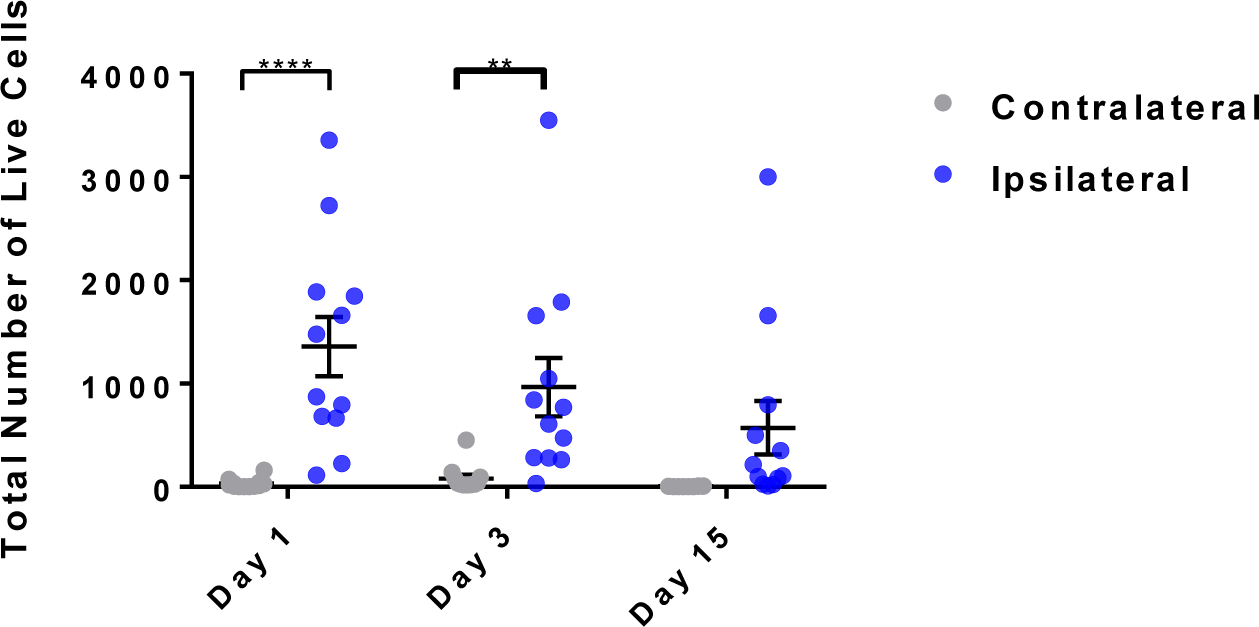
Neutrophil numbers in sciatic nerve peak at day 1 after partial sciatic nerve ligation. Plotted are the total number of live neutrophils (CD45+/Cd11b+/Ly6G+ singlets) isolated from sciatic nerve after partial sciatic nerve ligation at different time points. Each time, n = 12 mice per time point were processed (6 males & 6 females), shown here as separate data points with mean and SEM. Neutrophils were significantly upregulated one and three days after PSNL (two-way ANOVA, main effect of injury, F(1,66) = 33.47, p < 0.0001, significant at day 1 and day 3, p < 0.0001 and p = 0.0065, respectively).

**Supplementary Figure 8:**
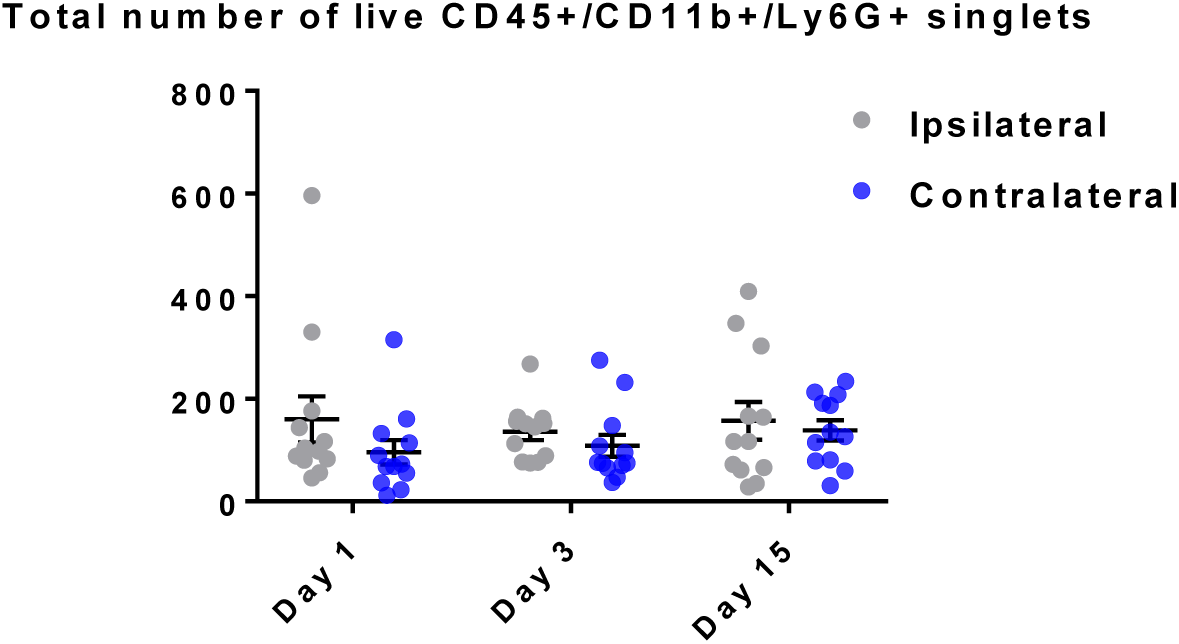
DRG contains a permanent CD45+/CD11b+/Ly6G+ population that does not vary with injury. Plotted are the total number of live neutrophils (CD45+/Cd11b+/Ly6G+ singlets) isolated from DRG after partial sciatic nerve ligation at different time points. Each time, n = 12 mice per time point were processed (6 males & 6 females); shown here as separate data points with mean and SEM. No significant differences in live cell numbers positive for these markers were detected in ipsilateral vs. contralateral DRG.

**Supplementary Figure 9:**
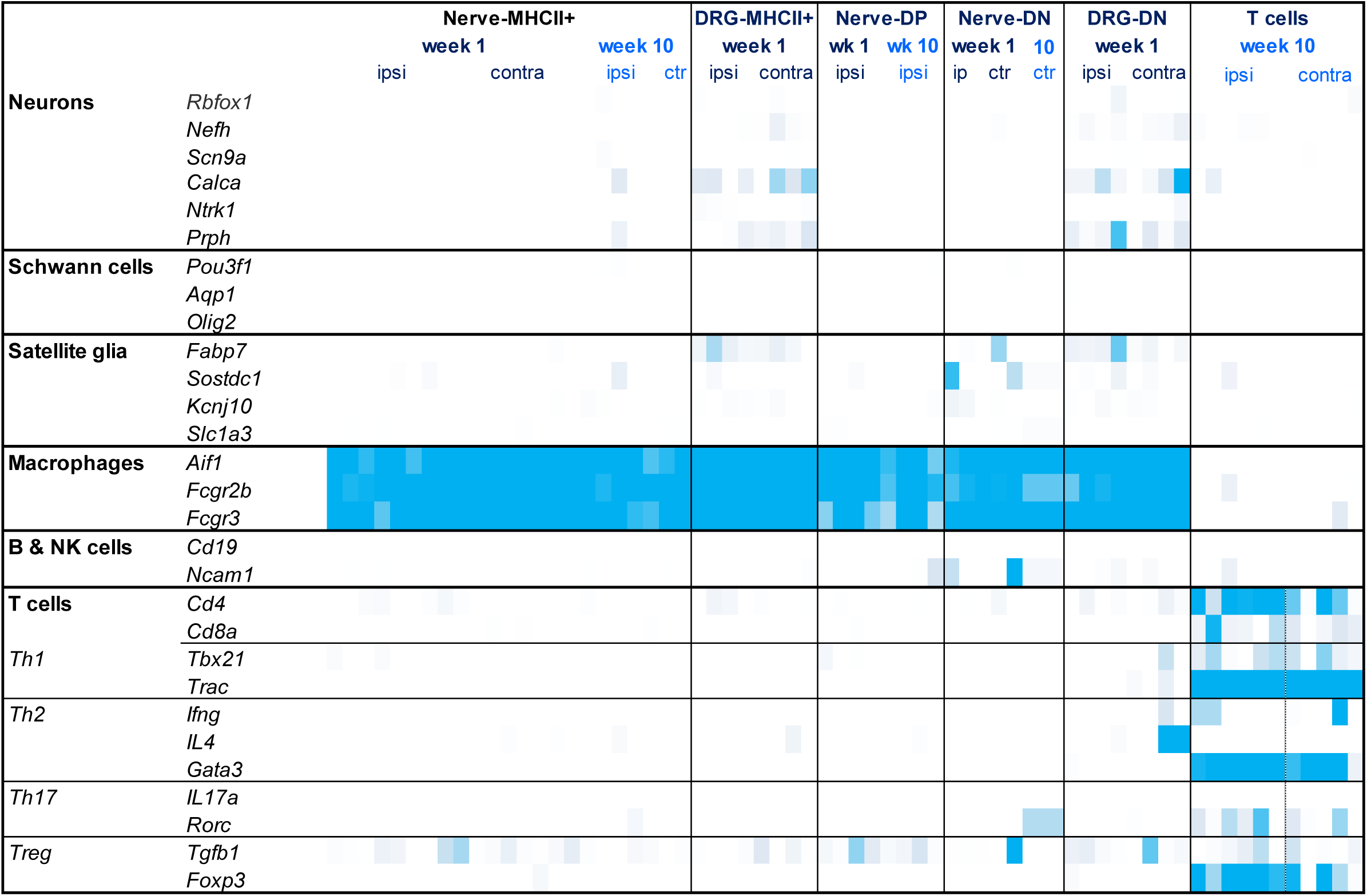
TPM expression levels for select marker genes to assess purity of sequenced samples. Myeloid and T cell samples from nerve were largely clean, with very little detectable contamination from neurons, Schwann cells and other lymphocytes. Samples from DRG had a small amount of neuronal and satellite glial cell contamination, likely caused by satellite glial “pulling down” neurons they envelope and being sorted on CD45+, which a small percentage of them is known to express.

**Supplementary Figure 10:**
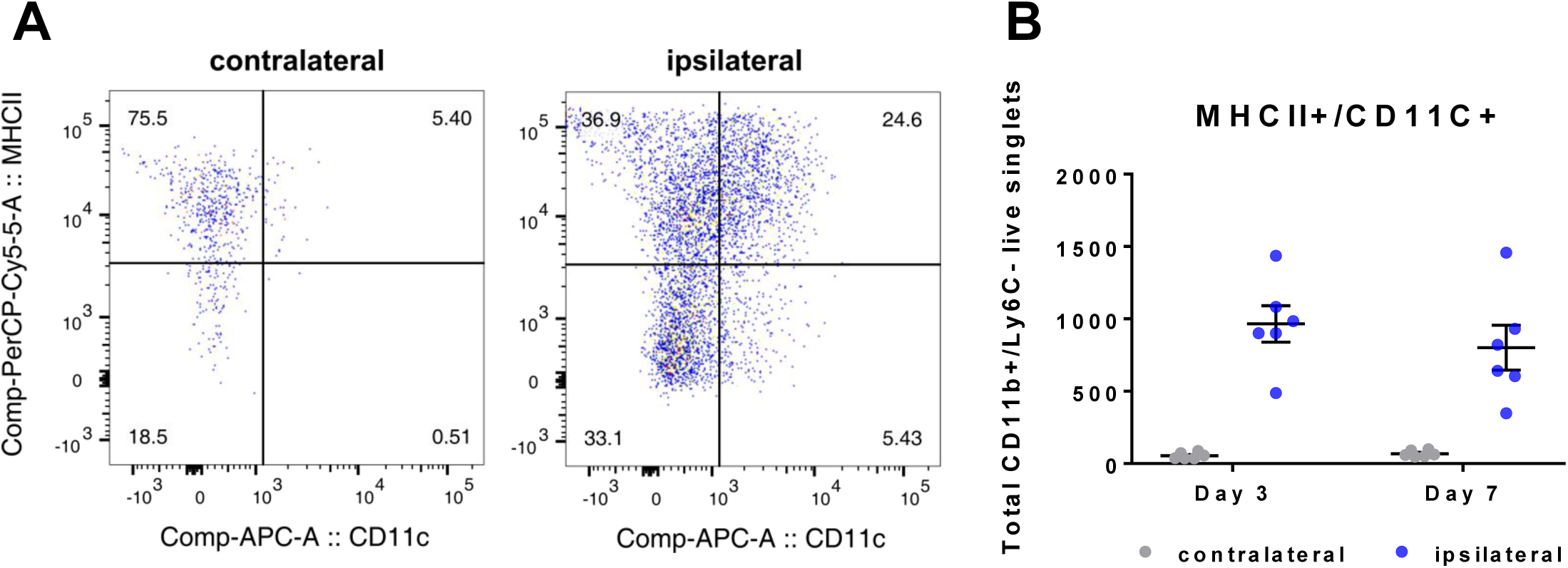
CD11c is upregulated at protein level in MHCII+/Ly6C-sciatic nerve macrophages seven days after PSNL. **A)** Representative dot plots of immune cells extracted from sciatic nerve seven days after PSNL. Live single cells were gated to be CD45+, CD11b+ and Ly6C-. CD11c was significantly upregulated in the MHCII+ population in ipsilateral samples. **B)** Quantification of the number of MHCII+/CD11c+ immune cells (gated on CD45+, CD11b+, Ly6C-live singlets) three and seven days after nerve injury: two-way ANOVA, F(1,20)=67.78, p< 0.0001. Each dot is an animal (n = 6, 3 male, 3 female) from which immune cells were obtained from ipsi- and contralateral nerves. Lines represent means and SEM.

**Supplementary Figure 11:**
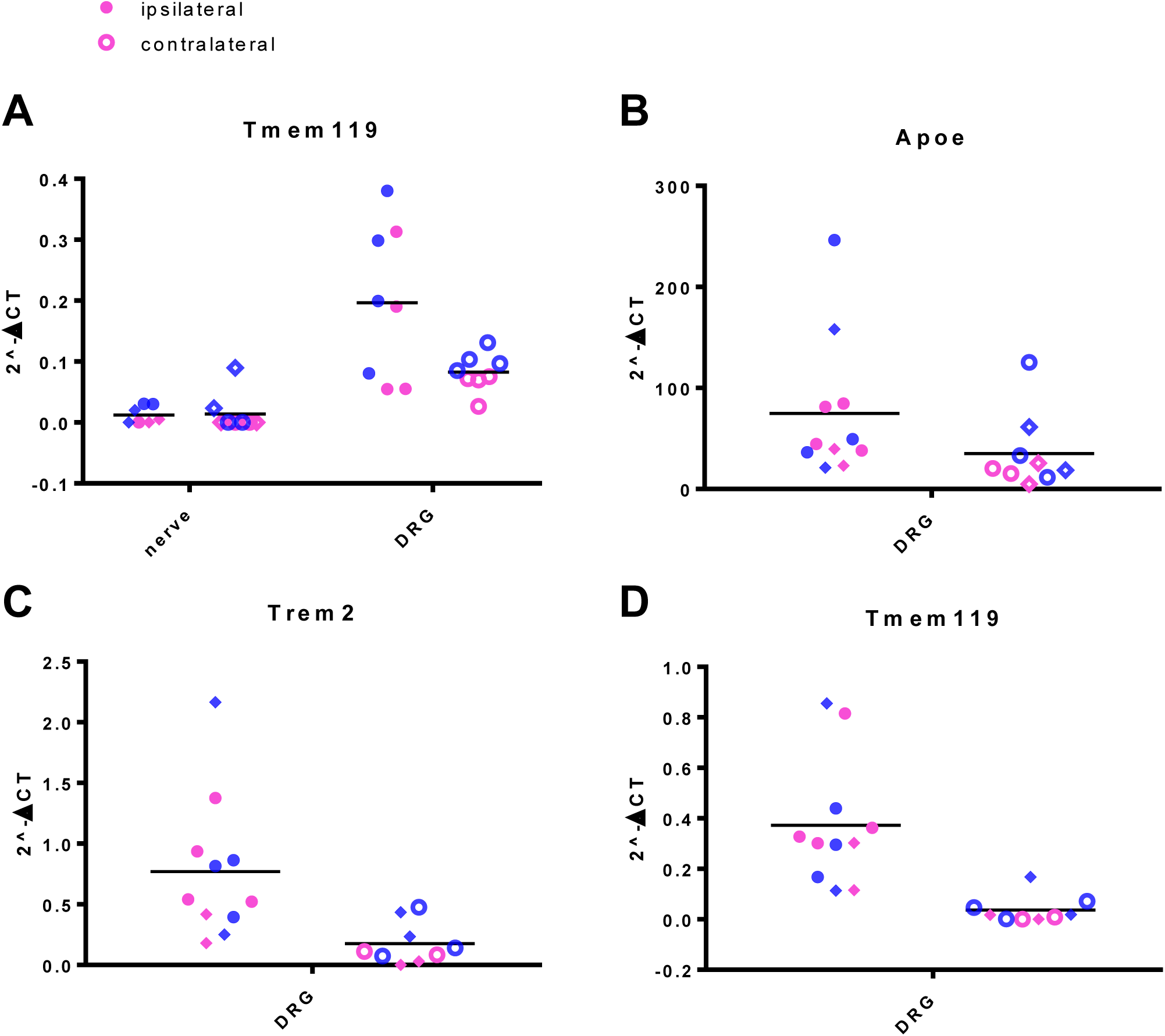
qRT-PCR confirming differences in expression of certain markers between nerve and DRG macrophages. **A)** qRT-PCR for *Tmem119* confirmed our sequencing results which indicated increased expression of this gene in MHCII+ macrophages from DRG compared to nerve: a two-way ANOVA revealed a significant main effect of *tissue* F(1, 27) = 26.4, p<0.0001 and significant interaction between *tissue x injury* F(1, 27) = 5.54, p = 0.0261. Each dot represents the 2^-ΔCT value of a biological replicate (blue = male, pink = female, n = 4). Samples drawn as diamonds were already included in our sequencing, so constitute technical replicates, while samples drawn as circles were independent biological replicates now additionally tested. **B-D)** qRT-PCR from MHCII-/Ly6C-macrophages extracted from ipsilateral and contralateral DRG (n = 11,9): two-way ANOVAs with *sex* and *injury* as factors revealed a significant main effect of *injury* for *Trem2* - F(1, 16) = 8.91, p = 0.0088, and *Tmem119* - F(1, 16) = 13.95, p = 0.0018, but not *Apoe*, in line with the p value in sleuth only passing false-discovery rate correction at p < 0.15 (Suppl. Table 5). Each dot represents the 2^-ΔCT value of a biological replicate (blue = male, pink = female). Samples drawn as diamonds were already included in our sequencing, so constitute technical replicates, while samples drawn as circles were independent biological replicates now additionally tested.

**Supplementary Figure 12:**
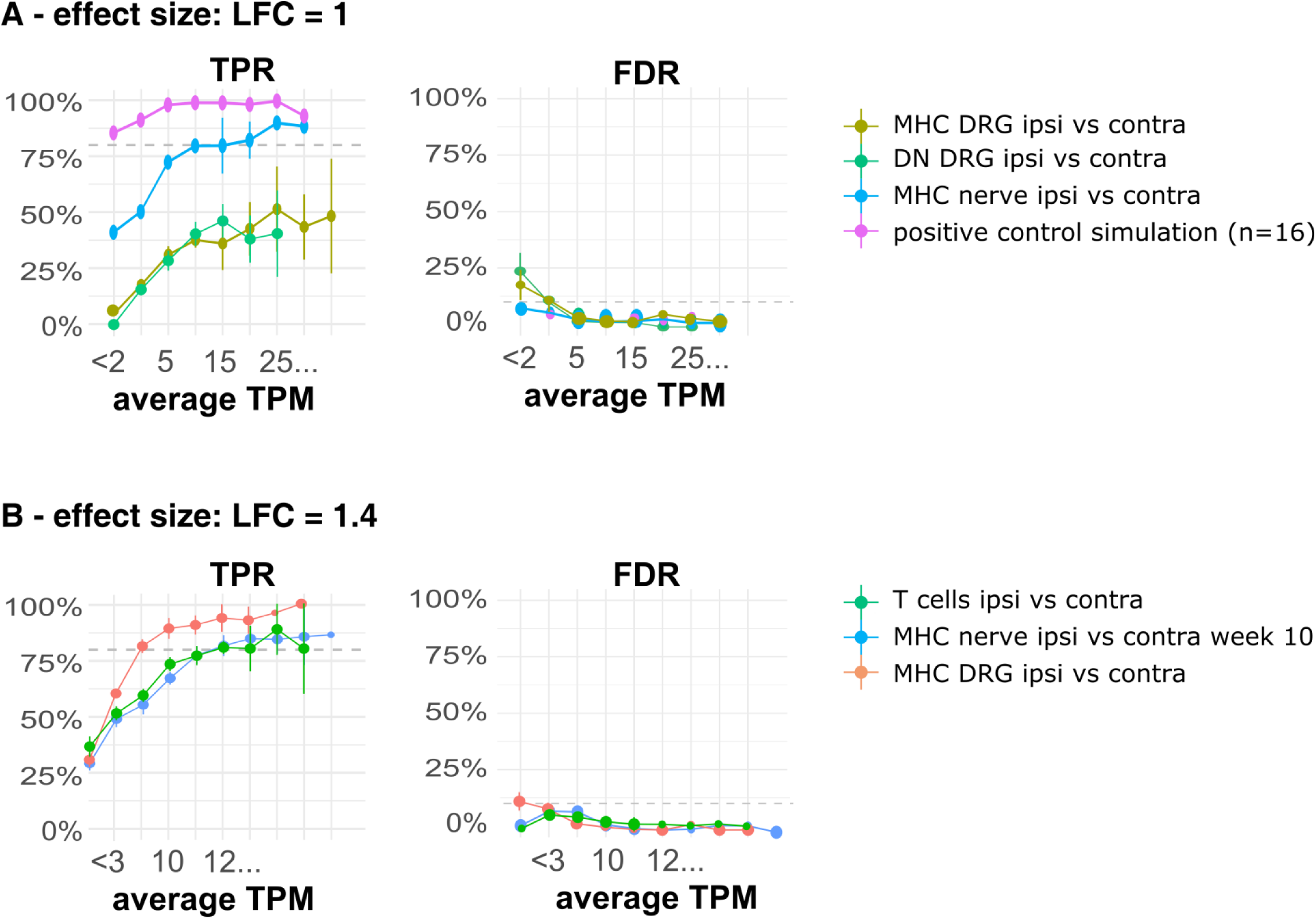
Results from powsimR package analyses, which employs our data as a seed to estimate the true positive effects and false discovery rates one would likely encounter with datasets of similar dispersion. **A)** The R package powsimR was fed raw count data from week 1 MHCII+ nerve and DRG, as well as MHCII-/Ly6C-(=DN) samples. The package estimates mean-dispersion relationships from these data and uses them as the basis for power analyses. TPR = true positive rate: the lines indicate the % chance to detect an effect size of log fold change (LFC) = 1, if one were to conduct differential expression analysis on data with similar properties, using the DESeq2 algorithm with n = 8 (the number of samples we had for MHCII nerve ipsi vs. contra comparisons), n = 4 (MHC DRG and DN DRG) and n = 16 (picked as a positive control). FDR = false discover rate: the lines indicate the chance of encountering false positives with the same simulation parameters. Data are split according to expression level: average TPM values were estimated from the mean log2 counts that are provided by powsimR. **B)** As in A, but this time the simulation was run to enable detection of a much larger effect size: log fold change (LFC) = 1.4. n numbers corresponding to week 10 T cell and MHCII comparisons and week 1 MHC DRG comparisons were used.

## Footnotes

1 Not in network graph, but significantly dysregulated – see Suppl. Table 9

